# Aptamer Conformational Dynamics Modulate Neurotransmitter Sensing in Nanopores

**DOI:** 10.1101/2023.03.10.532011

**Authors:** Annina Stuber, Ali Douaki, Julian Hengsteler, Denis Buckingham, Dmitry Momotenko, Denis Garoli, Nako Nakatsuka

**Author notes:** Annina Stuber and Ali Douaki contributed equally to this work.

## Abstract

Aptamers that undergo conformational changes upon small-molecule recognition have been shown to gate the ionic flux through nanopores by rearranging charge density within the aptamer-occluded orifice. However, mechanistic insight into such systems where biomolecular interactions are confined in nanoscale spaces, is limited. To understand the fundamental mechanisms that facilitate the detection of small-molecule analytes inside structure-switching aptamer-modified nanopores, we correlated experimental observations to theoretical models. We developed a dopamine aptamer-functionalized nanopore sensor with femtomolar detection limits and compared the sensing behavior with a serotonin sensor fabricated with the same methodology. When sensing these two neurotransmitters with comparable mass and equal charge, the sensors showed an opposite electronic behavior. This distinctive phenomenon was extensively studied using complementary experimental techniques such as quartz crystal microbalance with dissipation monitoring, in combination with theoretical assessment by the finite element method and molecular dynamic simulations. Taken together, our studies demonstrate that the sensing behavior of aptamer-modified nanopores in detecting specific small-molecule analytes correlates to the structure-switching mechanisms of individual aptamers. We believe that such investigations not only improve our understanding of the complex interactions occurring in confined nanoscale environments, but will also drive further innovations in biomimetic nanopore technologies.

## Introduction

Aptamer-integrated solid-state nanopores can serve as biomimetic systems that simulate how protein channels control ionic transport selectively in response to small-molecule binding. Aptamers are artificial, single-stranded oligonucleotides isolated through an iterative evolutionary method to interact specifically with analytes of interest.^1,2^ Advances in selection methodologies have enabled the discovery of aptamers targeting small molecules, ^2,3^ which are conventionally difficult targets with minimal available functional groups for recognition.^4–6^ Coupling such selective bioreceptors inside nanoscale pores with geometries that complement and confine the aptamer-target interactions, enables measurements that approach single-molecule sensitivities.^7–9^ Nanoscale pores can be prepared using different strategies.^10–13^ In particular, glass nanopipettes are one of the most convenient nanopore platforms due to the ease of fabrication *via* laser pulling, which can yield pore sizes ranging from a few to tens of nanometers with high reproducibility. ^14,15^

Herein, we have developed a specific and selective dopamine aptamer-modified nanopipette sensor with a nanoscale sensing area (10 nm nanopore) and femtomolar detection limits. Such nanoscale sensors for neurotransmitter detection approach synaptic dimensions, facilitating localized measurements with nanoscale spatial resolution. Implantable aptamer-based neural probes, while comparably sensitive and selective, are on the microscale.^16,17^ Similarly, other conventional methods such as microdialysis or fast scan cyclic voltammetry (FSCV) are limited in probe dimensions: the smallest probes to date are in the range of 100 *µ*m^18^ to hundreds of nm^19,20^ in diameter, respectively. Further, existing implantable neural probes or FSCV electrodes have exposed surfaces, which are prone to biofouling in biological environments.^21^ By confining the sensing area within an aptamer-filled nanopore that occludes nonspecific binders, we minimize surface biofouling and demonstrate the potential to measure dopamine in undiluted biofluids such as human serum.

Beyond developing highly sensitive nanoscale dopamine biosensors for neuroscience, our goal was to glean mechanistic insight into the function of small-molecule nanopore sensors driven by structure-switching. Detection of comparably sized and equally charged small molecules (dopamine and serotonin both carry a single positive charge per molecule under physiological conditions) using identical nanopore dimensions, allowed investigations into the influence of target-specific aptamer conformational dynamics on the measured sensor response. We observed a contrasting phenomenon when comparing the dopamine and serotonin nanopipette sensors using the same measurement system. The ionic current changed in opposite directions: a *decrease* was observed upon dopamine detection while an *increase* was recorded for serotonin sensing. While prior works implicated structure-switching aptamers to modulate the opening and closing of aptamer-modified nanopores, ^22–27^ in-depth characterizations of analyte-specific aptamer structural rearrangement in the context of nanopores has been lacking.

Thus, to understand the driving mechanisms for signal transduction in aptamer-modified nanopores, the conformational dynamics of the dopamine and serotonin aptamers were studied and compared experimentally and theoretically. The divergent directionality of structure switching for the two neurochemical aptamers on surfaces were corroborated by quartz crystal microbalance with dissipation monitoring (QCM-D). Further, molecular dynamics (MD) simulations, previously shown to predict oligonucleotide tertiary structures accurately,^28–32^ were performed to simulate the aptamers in their free *vs*. target-bound states, enabling visualization of the conformational dynamics of the serotonin and dopamine aptamers upon capturing their respective targets. Correlations between experimental and theoretical findings demonstrate the potential of MD simulations as an *in silico* tool to predict aptamer structure-switching mechanisms. Further, values of aptamer height change extracted from the simulations enabled calculations *via* the finite element method to correlate the electrochemical behavior of the sensor to aptamer-specific conformational dynamics inside the nanopore.

## Results/Discussion

### Specific Dopamine Sensing *via* Aptamer-Modified Nanopipettes

The current flux through the nanopore is measured upon application of a bias between the two Ag/AgCl quasi-reference electrodes: one inside the nanopipette and one in the bulk solution (Figure 1a). Dopamine-specific DNA aptamers were functionalized using sequential surface chemistry on the inner wall of quartz capillaries with *ca*. 10 nm orifices. The nanopore opening size was previously visualized using transmission electron microscopy.^27^ Further, finite element method simulations of the ionic flow through the nanopore corroborated this pore size - the experimental electrical resistance value of 367 MOhm, suggests an opening of 9.2 nm (for details on the simulations, see Supporting Information).

**Figure 1:**
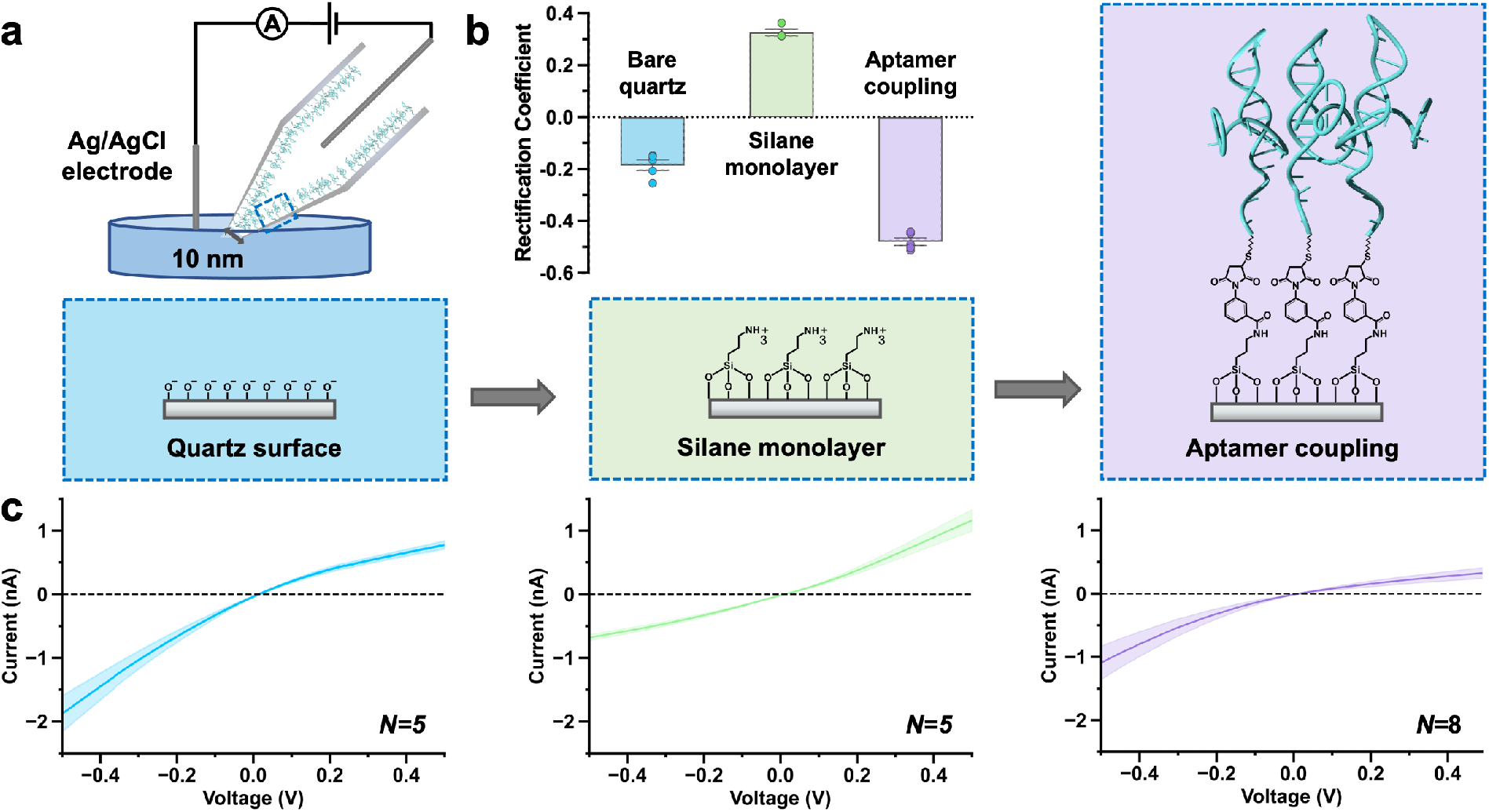
(a) Dopamine aptamer-modified nanopipette sensing schematic and functionalization protocol. (b) The rectification coefficient was calculated to obtain numerical values for the surface charge at each step of the functionalization protocol and to demonstrate aptamer assembly. (c) Dopamine aptamers functionalized through sequential surface chemistry can be tracked using the ion current rectification (ICR) effect manifested as asymmetric current *vs.* voltage curves. Negative charges on bare quartz nanopipettes (*N* =5) are inverted to positive upon assembly of aminosilanes (*N* =5). Coupling of negatively charged aptamers leads to higher rectification behavior (*N* =8). The solid line represents the average and the shaded area the standard error of the mean.

Ion transport through nanopipettes shows non-Ohmic behavior, where ionic current in one direction is favored due to the influence of an asymmetric electrical double layer in a confined area.^33^ This non-linear, diode-like effect observed in the current-voltage curves is called ion current rectification (ICR), which is influenced by the surface charge inside the nanopore when the pore size and geometry are kept constant.^34–36^ The rectification coefficient (*r*) can be extracted from the ICR by taking the logarithm of the ratio of the absolute values of the current measured at a positively applied voltage, by the current at the corresponding negative voltage:

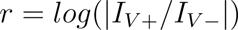

The *r* value changes from −0.2 *±* 0.03 for bare quartz surfaces due to deprotonated hydroxyl groups to +0.3 *±* 0.02 upon assembly of positively charged aminosilanes (Figure 1b). Upon covalent attachment of the dopamine aptamer, a higher negative charge (−0.5 *±* 0.03) compared to the bare quartz surface was observed due to the phosphate backbone of the DNA.

The ICR effect manifested as non-linear curvature in the current-voltage characteristics, enables tracking of the each step during the sequential surface chemistry for aptamer immobilization inside the nanopipette (Figure 1c). The negative ICR of a bare quartz surface implies that the ionic transport is favored at negative applied potentials. Upon aminosilane assembly, the curvature is inverted, implying favorable transport at positive applied potentials. Coupling of aptamers returns the ICR to a distinctly curved negative rectification. Tracking the ICR and *r* value is a route to ensure optimal starting nanopore sizes post fabrication and effective surface modification in subsequent steps to increase functional sensor yield.

Upon exposure of dopamine aptamer-modified nanopipettes to increasing concentrations of dopamine (with 10 % wt ascorbic acid to prevent dopamine oxidation) in undiluted (1x) PBS, the sensor showed a decrease in the current response from baseline (Figure 2a). Alternatively, nanopipettes modified with control sequences designed to have the same number and type of nucleotides as the specific aptamer but in a scrambled order to hinder dopamine recognition, showed minimal response to high dopamine concentrations in PBS (Figure 2b). The lack of response of the control sensor in the presence of dopamine and ascorbic acid, demonstrates that the ascorbic acid neither alters the ionic milieu nor binds to DNA nonspecifically. Sequence specificity was further demonstrated in the calibration curve where high amounts of dopamine resulted in negligible responses for the control sensor while dopamine sensors showed a concentration-dependent response (Figure 2c). The dopamine aptamer-modified nanopipettes were sensitive to dopamine amounts in the fM - pM concentration range with a limit of detection of 1 fM and saturation at 1 nM.

**Figure 2:**
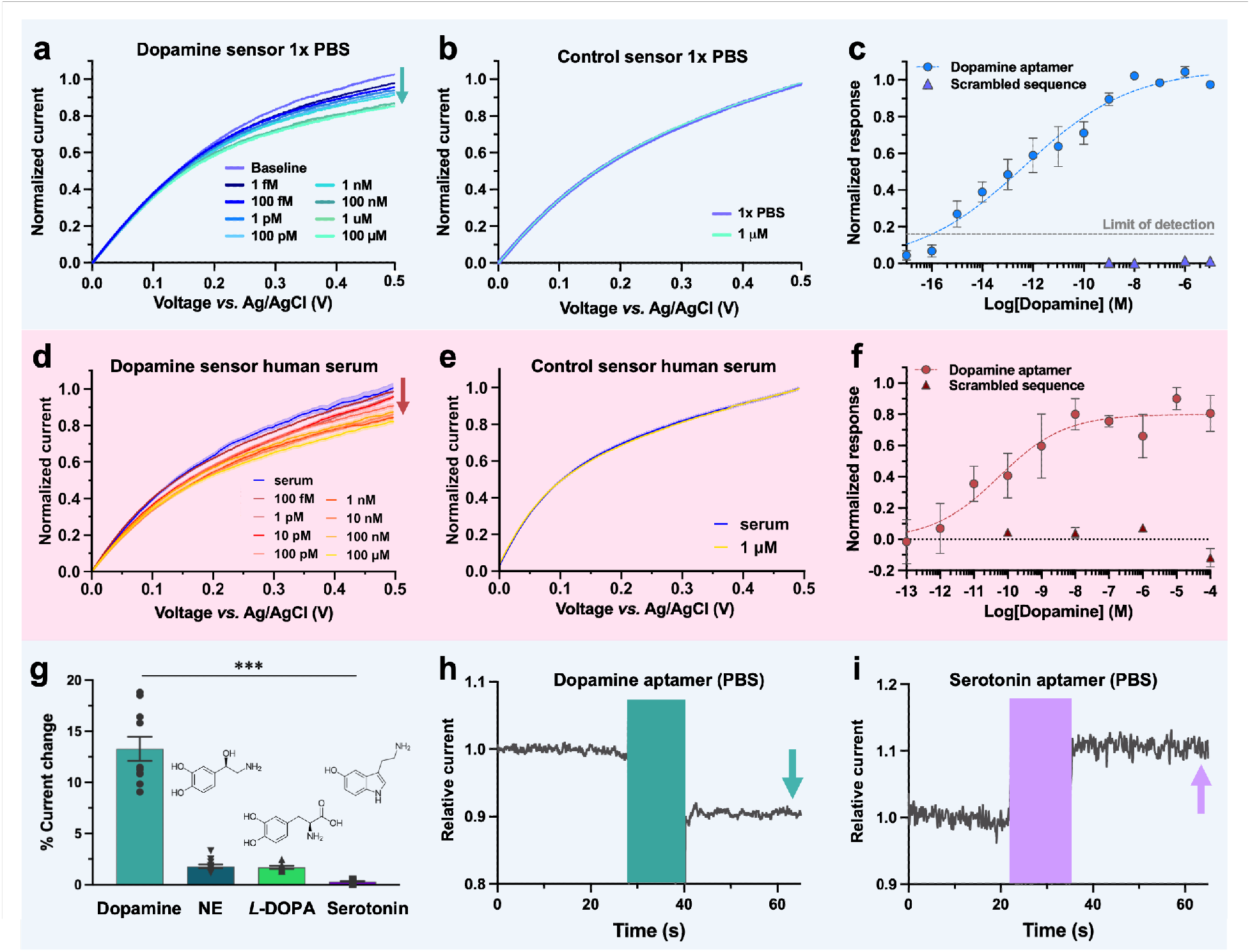
(a) Concentration-specific response of dopamine sensors in 1x phosphate buffered silane (PBS) with increasing dopamine amounts observed in cyclic voltammograms (CV). (b) The control sensor functionalized with scrambled DNA responded negligibly to high dopamine amounts (1 *µ*M). (c) Values extracted at +0.5 V from dopamine-specific CVs were used to plot calibration curves. Dopamine was detected in the 1 fM - 100 pM range by specific sensors (blue circles) while scrambled control sensors showed negligible responses (purple triangles). Each point is an average of *N* =5 CVs and for *N* =5 independent sensors). The blue dotted line is drawn to guide the eye but does not represent classical equilibrium binding. The limit of detection (signal at zero analyte concentration plus three times its standard deviation) is shown by the grey dotted line. (d) Comparable current decreases upon dopamine detection was observed in undiluted human serum. (e) Despite the increased complexity, the control sensor showed negligible changes upon dopamine exposure (1 *µ*M). (f) Calibration curves were constructed for both the specific (*N* =3) and control (*N* =2) sensors. As measurements were conducted in human serum that may already have basal levels of dopamine, the detection limit was not determined. (g) Dopamine sensors (13.3 *±* 1.2 % sensor response) demonstrate selectivity in neurobasal medium by differentiating structurally similar molecules such as norepinephrine (NE, 1.8 *±* 0.2 %) and *L*-3,4-dihydroxyphenylalanine (*L*-DOPA, 1.7 *±* 0.2 %) as well as analogously charged serotonin (0.3 *±* 0.05 %) with statistical significance for *N ≥* 10 sensors [one-way ANOVA: F(3,37)=107.7 p*<*0.0001]. (h) Real-time recording of the dopamine sensor in 1x PBS exposed to 100 *µ*M dopamine showed a current *decrease* while (i) serotonin aptamer-modified nanopipettes exposed to 100 *µ*M serotonin showed an *increas*^8^*e* in sensor response. Measured currents were normalized to baseline recordings in 1x PBS for comparative purposes.

The dopamine aptamer binding affinity (K*_d_*) has been reported as 150 nM.^37^ The observed non-linear behavior and detection limit orders of magnitude below the K*_d_* suggests that aptamer-modified nanopipettes are non-equilibrium sensors. Shift from conventional equilibrium behavior may arise from the conical geometry and nanoconfinement of the sensors. The frequency of target rebinding events may be higher at the most confined tip region compared to unbinding and release of targets occurring more prominently further away from the tip that tapers to larger volumes. Single molecules captured from a larger volume and concentrated at the nanopipette tip has been shown to enable fM detection limits.^38^ Inducing mass transport of charged molecules like dopamine through the nanopore by applied voltages, drives the analyte trajectory through an aptamer-filled column, thus augmenting molecular interactions. Macromolecular crowding has been shown to enhance the sensitivity through cooperative effects between electrolytes and polymers.^39^ ^40^

The orifice dimension and the resulting tightly packed aptamers occluding the nanopore is critical for sensitive biosensing. Dopamine could not be detected when the nanopore diameter was doubled to 20 nm (Figure S1), aligning with finite element models that correlate nanopore size with optimized sensor response (see Supporting Information). Limiting the size of the nanopore and ensuring aptamer confinement has further advantages for biosensing in complex biological media. The DNA aptamers pre-clog the pore to prevent nonspecific molecules from entering, which enabled measurements in undiluted human serum. Dopamine detection in human serum has important implications in neurodegenerative disorders such as Parkinson’s and Alzheimer’s.^41^ A current decrease of comparable magnitude to measurements in 1x PBS was observed for dopamine aptamer-modified nanopipette sensors with increasing amounts of dopamine in human serum (Figure 2d). Alternatively, control sensors with scrambled DNA, showed negligible changes in ionic behavior in serum (Figure 2e). Control sensors with comparable chemical signatures to aptamer sensors, serve as ideal references by differentiating signal changes due to the presence of dopamine *vs.* environmental perturbations or nonspecific binding. Such differential measurements are especially important in complex biological environments.^42^ The dopamine aptamer-modified nanopipettes detected dopamine in a concentration-dependent manner in human serum, while the control sensor showed negligible changes (Figure 2f).

As the human serum may have basal dopamine present, the detection limit was not determined but rather, the feasibility of sensing in complex environments was demonstrated. The quantification of unknown dopamine levels in clinical samples will necessitate standard addition measurements, a technique that introduces known analyte amounts to diluted samples to mitigate matrix effects. Implementation of differential measurements where the specific sensor is deployed in parallel to a control sensor, will further account for matrix-related effects. Herein, to demonstrate the viability of our sensor application in undiluted complex media, selectivity tests were conducted in neurobasal medium. This biofluid is devoid of basal dopamine while containing various nonspecific amino acids and proteins that support neural cultures *in vitro*. Dopamine sensors differentiated structurally similar molecules including norepinephrine (NE) and *L*-3,4-dihydroxyphenylalanine (*L*-DOPA) as well as serotonin with statistical significance in this complex medium (Figure 2g). Such selectivity in the presence of interferents renders aptamer-based sensors advantageous compared to antibody-based methods that suffer from cross reactivity and voltammetric methods that have challenges in distinguishing dopamine analogs with overlapping oxidation signals. ^43^

Translational strategies for these sensors are contingent on the specific deployment environment. Concentration ranges for basal and stimulation-evoked dopamine in the brain range from low nanomolar to micromolar levels^44–46^ while serum dopamine levels have been reported in the picomolar to high nanomolar levels.^47–49^ Consequently, calibration of the sensing regime is critical for use in specific applications. Sensor sensitivity can be tuned from lower to higher concentrations by modifying aptamer sequences^50–52^ or by adjusting the aptamer density on sensor surfaces.^37^ Therefore, a sensor capable of detecting femtomolar concentrations of dopamine in PBS offers flexibility in adjusting the detection range according to the translational needs.

However, to modulate sensor characteristics effectively, an understanding of the detection mechanism is critical. For the dopamine aptamer-modified nanopipettes, a consistent *decrease* in the current response from baseline was observed upon specific recognition of dopamine in undiluted PBS, serum, and neurobasal medium. In contrast, prior investigations with serotonin aptamer-modified nanopipettes demonstrated an *increase* in the current response from baseline with higher amounts of serotonin in the same environments.^27,53^ We show a side-by-side comparison of the real-time response from the dopamine (Figure 2h) and serotonin aptamer-modified nanopipettes (Figure 2i) in PBS upon exposure to their respective targets to highlight this divergent current response.

We hypothesized that the opposite directionality observed in the nanopipette sensors is driven by the divergent structure-switching of aptamers inside the confined nanopore. The *r* values for the two sensors were extracted to enable direct comparisons of the surface charge at the walls of the nanopipette in an aptamer-specific manner (Figure S2). ^54^ For the dopamine aptamer-modified sensor, the *r* value becomes increasingly negative, while the *r* value of the serotonin aptamer-modified sensor increases in value upon serotonin capture. These results indicate that dopamine *vs.* serotonin aptamers alter the surface charge density in opposite directions upon target recognition within the nanopore.

### Tracking Aptamer Conformations *via* Quartz Crystal Microbalance

To corroborate our findings concerning the directionality of current change upon target recognition with a complementary methodology, QCM-D was employed. While measuring the binding of molecules with low molecular weights directly using QCM-D is challenging, ^4^ aptamer conformational dynamics that modify the hydration layer at the surface of the sensor can be harnessed to amplify the signal of small-molecule binding.^55^ The restricted degrees of freedom, mimicking the dynamics inside the nanopore sensors, gives QCM-D qualitative insight into both the directionality and magnitude of conformational dynamics for aptamers tethered to surfaces. Dopamine aptamer assembly on the QCM-D substrate was confirmed by a decrease in frequency of 23.7 Hz (Figure 3a) and an increase in dissipation of 0.9 x 10*^−^*^6^ (Figure S3a) indicative of an assembled rigid monolayer with a height of 4.1 nm, calculated *via* the Sauerbrey equation (Figure 3b).^56,57^ Rinsing the surface post-assembly resulted in negligible changes in frequency, confirming the covalent assembly of thiolated aptamers on the surface of gold QCM chips. Upon exposure to dopamine, a frequency increase of *∼*3 Hz was observed (Figure 3c). This frequency increase translates to a decrease in aptamer height of *∼*0.6 nm (Figure 3d), which is indicative of the aptamers adopting more compact secondary structures upon dopamine binding, leading to loss of water molecules from the aptamer monolayer.^55^ The return to original baseline upon rinsing with buffer, demonstrates the reversible unbinding of captured dopamine.

**Figure 3:**
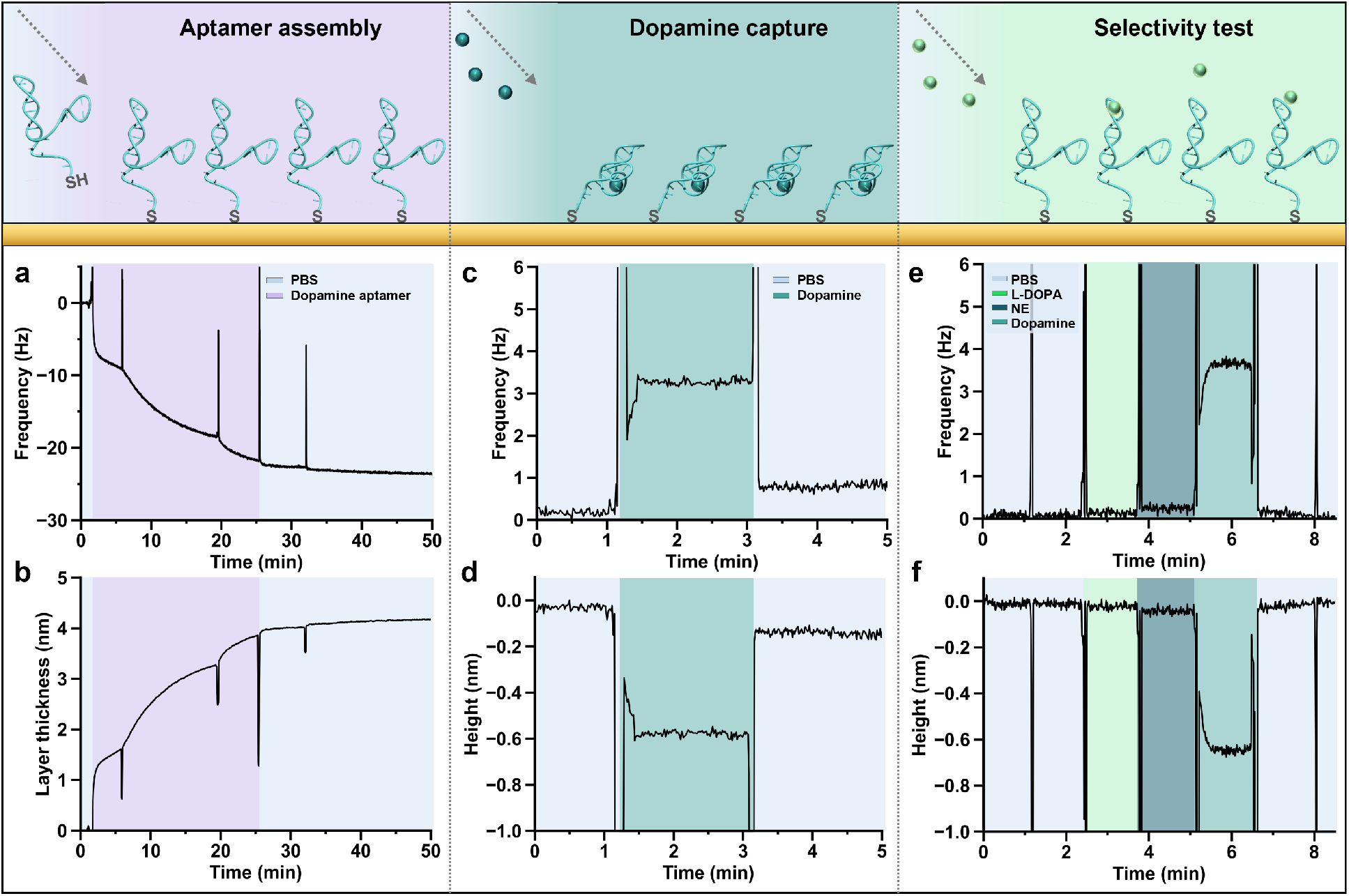
(a) Dopamine aptamer assembly in phosphate buffered saline (PBS) results in a frequency decrease of 23.7 Hz (b) The Sauerbrey equation calculates an assembled aptamer monolayer of 4.1 nm. (c) Exposure of aptamer-modified substrates to 100 *µ*M dopamine led to a reversible increase in frequency of 3.3 Hz. (d) This frequency change translates to a 0.6 nm compression in the aptamer layer upon dopamine recognition based on the Sauerbrey equation. (e) Neither injection of PBS nor 100 *µ*M of nonspecific molecules (*L*-Dopa and Norepinephrine (NE)) showed baseline changes while subsequent addition of dopamine exhibited the characteristic increase in frequency. (f) Despite prior exposure to nonspecific molecules, the dopamine aptamer compresses upon target recognition.

We demonstrated the reproducibility of this binding and unbinding behavior three repeated times on a single QCM-D sensor (Figure S3b) and over *N* =6 measurements conducted with three different chips (Figure S3c). When repeating this procedure on a chip functionalized with the scrambled control DNA, negligible change in frequency was observed upon exposure to equal amounts of dopamine, indicating target-specific structural rearrangement (Figure S3d). Further, injection of the PBS buffer as well as incubation of structurally similar molecules (*L*-DOPA and norepinephrine) resulted in a stable baseline (Figure 3e,f). The subsequent incubation of dopamine, resulted in a frequency increase of comparable magnitude when the aptamer-modified chip was tested solely with dopamine. These experiments are indicative of negligible interference from exposure to nonspecific molecules; structure switching of the dopamine aptamer occurs only in the presence of the specific target.

The compression of the dopamine aptamer layer upon target recognition observed at the surface of the QCM-D sensor, is in the opposite direction to what was observed for the serotonin aptamers upon binding serotonin (1.2 nm elongation of aptamer backbone upon target capture).^27^ These surface-based experimental results supported our hypothesis that the distinctive conformational dynamics of individual small-molecule aptamers, influence the direction and magnitude of the measured current through the nanopore. While dopamine concentration-dependent frequency changes were observed (Figures S4e,f), QCM-D is an ensemble method that monitors the dynamics of aptamer monolayers, which may not represent the behavior of individual aptamers within the nanoscale tip. To interrogate single molecule aptamer structure-switching mechanisms, molecular dynamics (MD) simulations were conducted.

### Simulating Target-Specific Aptamer Conformational Changes

The structure-switching dynamics of the dopamine and serotonin aptamers upon interactions with respective targets were conducted with a constraint on the 5’ end (typically modified with thiol groups for surface attachment to sensors) to mimic covalently-tethered states with reduced degrees of molecular freedom. The conformational stability of the aptamers can be determined by the deviations in the root mean square deviation (RMSD) within the simulation time interval - smaller deviations indicate higher structural stability. The conformation dynamics were simulated for an interval of 100 ns, which guaranteed observation of the system in a stable state based on monitoring the RMSD (Figure S4).

The binding of the aptamer with a specific neurotransmitter target was monitored by MD simulations over the same time interval. Analysis of the aptamer binding phenomena elucidates the interaction of individual nucleobases involved in target recognition and the resulting intermolecular interactions. Analysis revealed that the dopamine aptamer—target complex was maintained *via* four stable hydrogen bonds, electrostatic interactions (Pi-Anion, Pi-Pi-T-shaped, salt-bridge), and hydrophobic interactions (Figure S5a). Specifically, hydrogen bonds formed between the hydrogens of dopamine and the aptamer nucleobases A28, A30, and G33 at distances of 3.12 Å, 4.42 Å, and 3.08 Å, respectively, constitute the binding pocket (Figure 4a). Moreover, the primary amine (protonated at physiological pH) of the dopamine interacted electrostatically with the nucleobases G33 (Pi-cation), A35 (salt-bridge), and with A27 (Pi-anion) (Figure 4b). Further, hydrophobic Pi-Pi T-shaped interactions between G29 and the aromatic moiety of dopamine can be observed.

**Figure 4:**
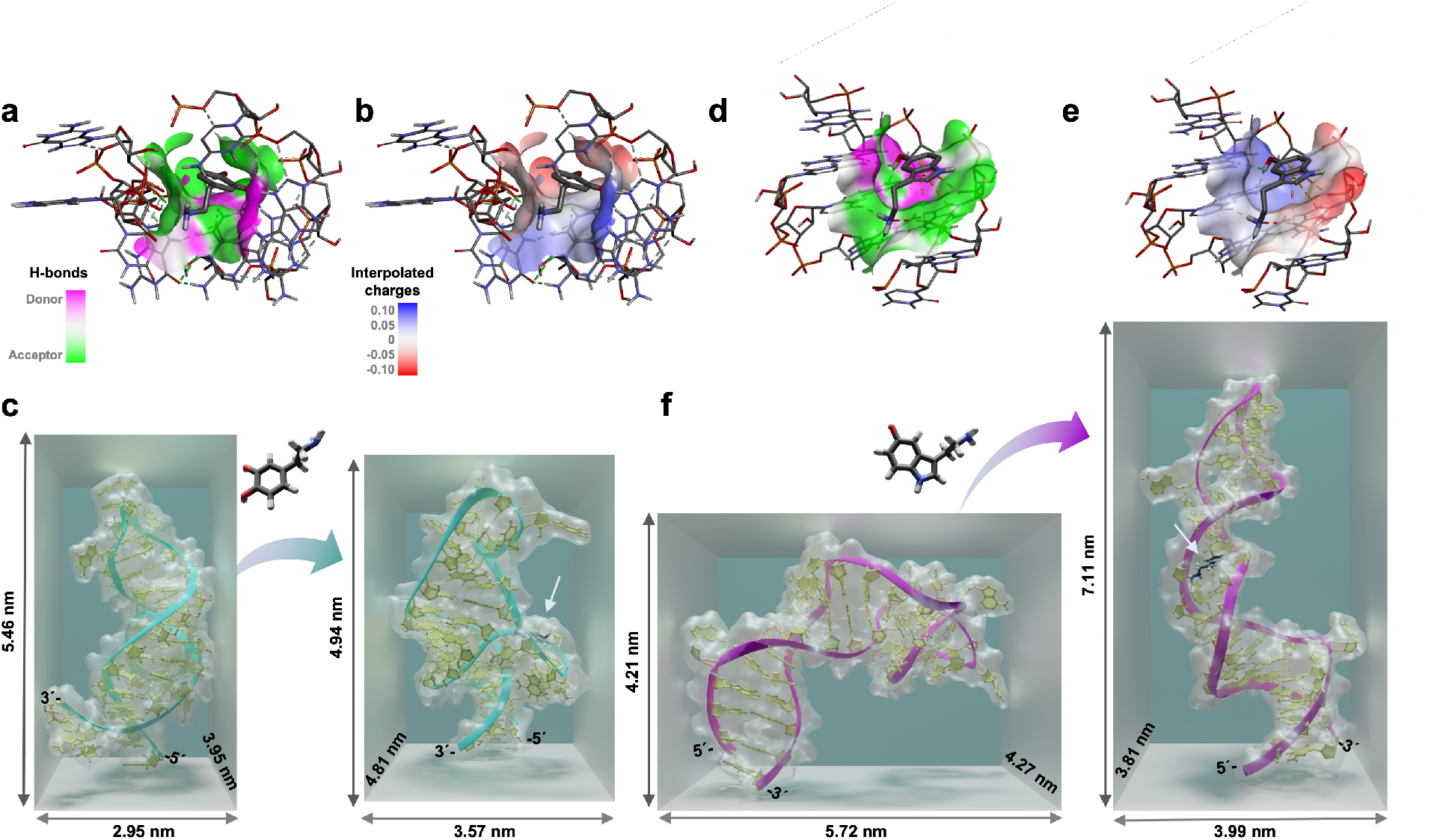
(a) Hydrogen bonding and (b) interpolated charges (blue = more positive, and red = more negative) interactions between the dopamine aptamer and dopamine molecule that constitutes the binding pocket. (c) Extracted 3-D conformations of the dopamine aptamer in the absence (left) and presence (right) of dopamine obtained from 100 ns molecular dynamic (MD) simulations (d) The hydrogen bonding and (e) interpolated charges between the serotonin aptamer and serotonin molecule. (f) 3-D structure of serotonin aptamer in the absence (left) and presence (right) of serotonin obtained from 100 ns simulations. In both bound-states for the dopamine and serotonin aptamers, an arrow is added as a guide to the eye to visualize the binding location of the respective analyte. All aptamer sequences were confined at the 5’ end to mimic surface tethering. To improve the visibility, all water molecules and ions inside the simulation box were removed.

From these intermolecular interactions, where the bases A28, A30, G33, A35, A27, and G29 interact strongly with the dopamine molecule, it can be inferred that the binding takes place inside the asymmetric interior loop of the dopamine aptamer. The 3-D structure of dopamine aptamer in the absence and presence of dopamine was extracted at the end of the MD simulation (Figure 4c). According to this visualization, the dopamine binding site is located at the distal stem-loop (or hairpin loop). Dopamine binding led to a conformational change where the aptamer-target complex compresses by 0.6 nm, comparable to the findings in QCM-D (Figure 3d).

Target-specific interactions for the serotonin aptamer was also investigated *via* MD simulation to interrogate the opposite behaviors observed in the serotonin *vs.* dopamine sensors. The aptamer-serotonin complex consists of four stable hydrogen bonds and electrostatic interactions (Pi-Anion, favorable acceptor-acceptor, and van der Waals) (Figure S5b). Explicitly, the binding pocket was composed of hydrogen bonds formed between the hydrogens of the serotonin and the G19, G20, T26, and G27 aptamer nucleotides with a distance of 2.12 Å, 2.59 Å, 1.89 Å, and 2.79 Å, respectively (Figure 4d). Moreover, electrostatic interactions between the nucleotide A17 and the 6-membered ring of the serotonin, and between the nucleotide G18 and the primary amine of serotonin were observed (Figure 4e and S6b). The 3-D structures of the free serotonin aptamer and the target-bound state with the serotonin molecule in the distal stem loop were extracted (Figure 4f). The simulation indicates that the serotonin aptamer backbone elongates upon serotonin binding, in agreement with previous experimental analyses.^37^

The MD simulations enabled visualization of the 3-D structures of individual dopamine and serotonin aptamers. Further, the binding pockets to respective targets were identified based on the energetics of intermolecular interactions. While methods such as circular dichroism spectroscopy and Förster resonance energy transfer (FRET) enable tracking aptamer structural rearrangements upon target binding,^6^ such ensemble measurements are limited. Circular dichroism can infer the formation of specific structural motifs but cannot indicate the location of rearrangement. The change in distance of certain bases in the aptamer backbone can be monitored locally *via* FRET. However, mapping the entire structural dynamics from FRET is challenging; the attachment of fluorophores in certain locations of the DNA backbone can interfere with structure switching and target recognition. To this point, MD simulations of aptamers and their respective targets contribute structural information at the single molecule level.

### Modeling Aptamer Conformational Changes in Nanopores

Experimental results by QCM-D and theoretical analysis by MD simulations both indicate that the conformational changes of aptamer molecules modulate the measured ion current changes in nanopipette sensors. To connect our findings on the aptamer structure-switching dynamics to the ionic current modulation through nanopores, a finite element model was designed. The model consisted of a solid nanopore sensor (geometry determined from electron microscopy images) modified with an ion-permeable charged layer with variable thickness, representing the aptamers. We aimed to interrogate the effect of altering the aptamer layer charge density on the rectifying behavior of the nanopore sensors. We expected the signal transduction through the nanopore to be determined by two phenomena: (i) variation of the stored charge density in the layer of the aptamer molecules at the nanopore walls, and (ii) changes in the aptamer layer permeability for ions carrying the current. The former occurs as the expansion/contraction of the aptamer layer can cause a proportional decrease/increase in the number of charges per unit volume respectively, even when assuming no overall change in the total number of charges.

Both the QCM-D and MD simulations indicated that the dopamine aptamer layer compresses upon target recognition. The values extracted from the simulation for the free aptamer (5.5 nm) and dopamine-bound aptamer (4.9 nm) were used to approximate the change in the charge density in the dopamine aptamer layer. An increase in the charge density by *∼*11 % (proportional to the layer thickness change), was used as a parameter in the finite element model (Supporting Information SI-5). The simulation for the dopamine aptamer-modified nanopipettes indicated that the contraction of the charged aptamer layer on nanopore walls, resulted in stronger rectification and a slight decrease of the ion current at positive voltages. This trend matches what was observed in the experiments upon sensor exposure to dopamine.

However, the charge density variation alone did not account for the experimentally observed changes of the current magnitude. For the dopamine sensors, we hypothesize that the contraction of the dopamine aptamer layer leads to a more compact and hence less permeable medium, through which ions are transported with a smaller diffusion constant.^58^ When a proportional reduction of the diffusion coefficient of ions is introduced to the charged layer of aptamers in the model, a further reduction of the ion current through the nanopipette is observed ((Supporting Information SI-5). In this case, the total change of the ionic current for the free dopamine aptamer *vs*. dopamine aptamer-target complex approaches the experimentally observed 11 % current decrease for 1 nM analyte. However, the difference in curve shapes observed in the model compared to the experimental I-V characteristics, suggests a qualitative agreement rather than quantitative comparison.

The finite element model appears to be generalizable for different aptamers when the sequence-specific conformational dynamics is well understood. While the behavior and the measured signals follow an opposing trend for the dopamine *vs*. serotonin aptamers, the mechanism behind the signal transduction bears close similarities. Conformational change upon analyte binding appears to cause a contraction (dopamine) or expansion (serotonin) of the charged aptamer layer, accompanied by a change in mass transport (altered diffusion constant) through this medium. Interestingly, these factors are most pronounced for both negatively charged aptamers only in the positive side of the I-V curve. Alternatively, at negative biases, the effect of changing the thickness of the charged layer and the influence of altering the ion diffusivity tends to cancel each other out, resulting in a lack of observable trend for sensing.

When taking these effects into account, the model also enables determination of the optimal aptamer/pore configuration for sensing (see Supporting Information SI-5). Two factors play major roles: (i) the initial thickness of the aptamer layer (before addition of target analyte) and (ii) the total change of the aptamer layer height upon target addition.

According to these calculations, the optimal initial aptamer layer should be around 5 nm (for a nanopore with 9.2 nm opening). Our dopamine aptamer layer thickness of *∼*5.5 nm determined through MD simulations (Fig. 4c) is nearly fully optimal for sensing with our *ca.* 10 nm nanopores. Concerning the conformational change upon target recognition, the nanopore is highly sensitive to the 3-D change in the aptamer geometry, suggesting that the larger the aptamer structure switching (*i.e.,* layer thickness), the larger the influence on the current through the nanopore. Such a model enables predictions of optimal aptamer-nanopore confinement parameters based on magnitude of structure switching *vs.* nanopore size.

In addition to finite element modeling, we calculated the Dukhin number (*Du*), which identifies the ratio of the surface *vs*. bulk conductivity occurring within the aptamer-modified nanopore.^59^ The approximated *Du* value of 8.4 and 4.0 for the dopamine and serotonin aptamer-functionalized nanopores, respectively (calculations can be found in Supporting Information, section SI-7), implies that surface conductivity dominates *vs.* the bulk (Figure 5). The measured current through the nanopore is primarily driven by the ionic flux that resides within the Debye layer at the walls of the nanopore (corresponding to *∼*0.7 nm in 1x PBS).^59^

**Figure 5:**
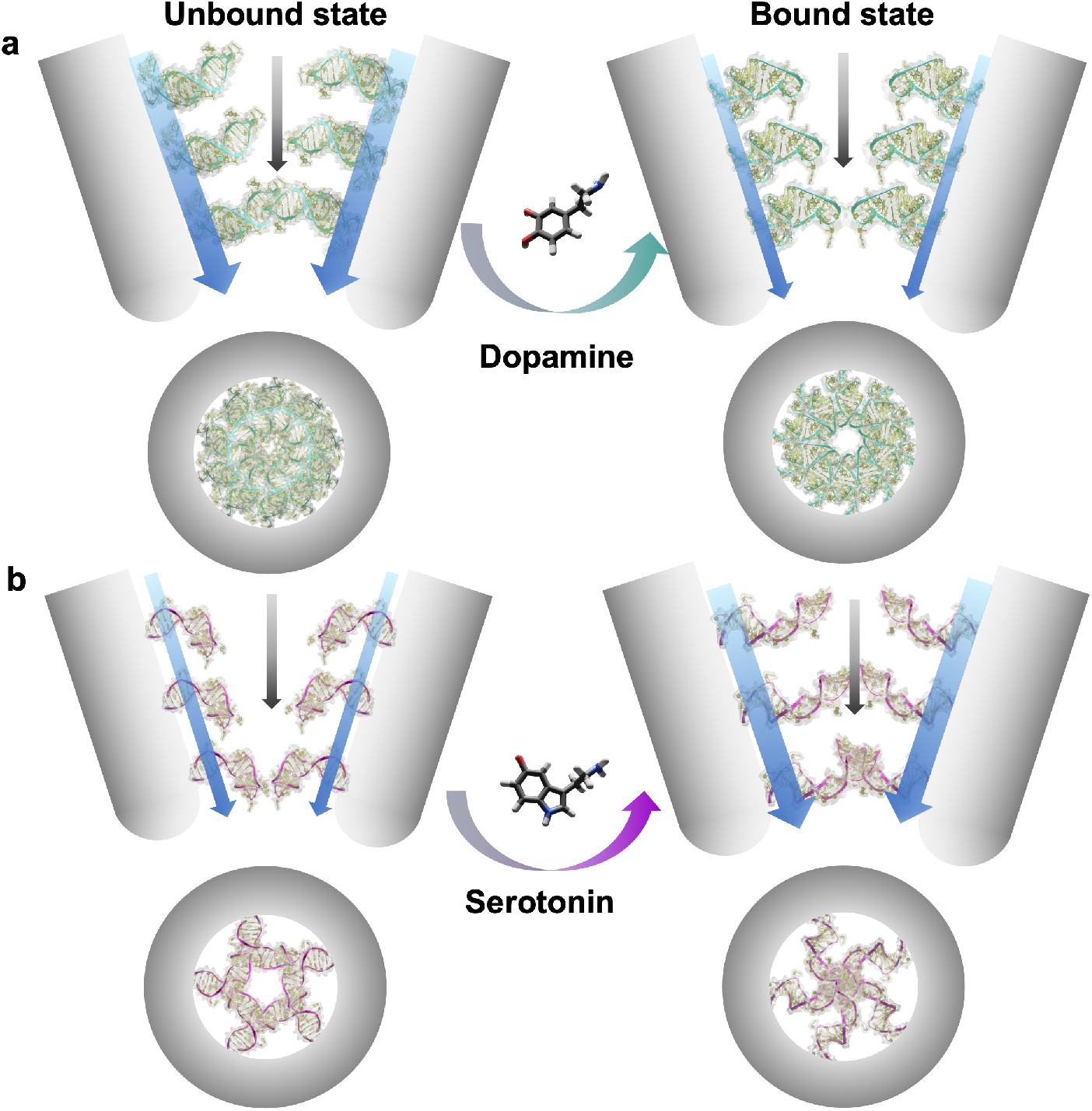
Schematic representation of ionic flux through dopamine and serotonin aptamer-functionalized nanopipettes. (a) Dopamine and (b) serotonin aptamers are shown as immobilized on the walls of the nanopipette from a side-view (top) or when looking inside the 10 nm orifice (bottom). The maximal number of aptamers bound at the tip was estimated by using the minimal required surface corresponding to the bottom rectangle extracted from the molecular dynamic simulation of the free aptamer. The unbound aptamer state is shown on the left and the target-bound state on the right. The respective targets trigger conformational changes altering the charge density and DNA layer thickness inside of the pore. The blue arrows represent the cationic flux along the walls of the nanopipettes, while the black arrows represent the cationic flux of the bulk of the nanopipette. The width of the arrows is used to represent magnitude of the flux under the different DNA conformations.

Thus, when the dopamine aptamers contract upon target recognition, this conformational change will influence the ionic conductance along the nanopore walls (represented by the blue arrows in Figure 5), while the bulk ionic flux through the nanopore remains unchanged (black arrows in Figure 5). Moreover, the mobility of solution ions within the DNA layer is contingent on the conformational state of the aptamers. Tighter bundles of DNA reduce ionic permeability. ^58,60^ Thus, a contracted layer of dopamine aptamers will lead to decreased ionic flux and *vice versa* for the serotonin aptamers that expand upon target capture.

An alternative way to interrogate the influence of the aptamer conformational dynamics on the ionic flux through the nanopore, is by calculating the different factors that contribute towards the change in current. The specific conformation of the aptamers influences the charge density of mobile counterions in the nanopore that neutralize the negative DNA backbone. Thus, the experimentally measured current response has contributions from the current excluded by the volume of the 3-D DNA (*I_excluded_*) and the current that flows through the assembled aptamer layer (I*_aptamer_*). These calculations support our prior experimental and theoretical findings (Supporting Information SI-8).^60^ By interrogating aptamer structure-switching dynamics extensively through different models and connecting these findings to the observed sensor response, we have improved our understanding of the distinctive behavior of aptamer-modified nanopore sensors.

### Conclusions

In this work, we characterized a nanoscale dopamine sensor with femtomolar detection limits in buffer conditions and retains functionality in complex biofluids such as human serum and neurobasal medium. The integration of nanopipette sensors as probes in various established platforms such as patch clamp setups and scanning probe systems, holds the promise for fast adaptation for diverse applications at the intersection of biological systems such as cell and tissue cultures. Moreover, we anticipate advancing small-molecule sensing in clinical samples by leveraging the advantages of cost-effective fabrication (*<*$1 per aptamer-modified nanopipette sensor) and real-time sensing capabilities. Our approach represents an advancement over existing detection systems, which require analyte separation and sample pretreatment. However, it is important to note that for clinical assessments, benchmarking against established gold standard methods will be imperative.

In parallel to the development of nanoscale dopamine sensors, we investigated the fundamental mechanisms governing the modulation of ionic flux through nanopores. In particular, our goal was to understand the opposite directionality of the measured current when comparing aptamer-modified nanopipette sensors targeting two neurotransmitters of comparable mass and equal charge: dopamine and serotonin. Through QCM-D, a complementary surface-based piezoelectric sensor, we confirmed divergent aptamer conformational dynamics upon target binding.

To corroborate these ensemble measurements from QCM-D with simulations conducted at the single molecule level, MD simulation was conducted. Correlations between the theoretical and experimental findings validated MD simulation as a potential layer of control for establishing design rules for aptamer-based nanopore sensors. For example, visualization of the aptamer-target interactions may provide insight into how to optimize aptamer surface densities to minimize steric hindrance or what size nanopores to employ based on magnitude of structure switching. We envision that integration of MD simulations within the aptamer selection process, could create an intermediate feedback loop to generate aptamers endowed with optimal structure-switching capabilities, which would accelerate the development of next-generation aptamer-modified sensors. Further, values extracted from the simulations can be incorporated into finite element models to understand the combined influence of changes in the charge density and ion permeability of the aptamer layer. Such investigations improve our understanding of the complex interactions occurring in confined nanoscale environments.

The central role of aptamer conformational change within nanoscale confinement as the driving mechanism for small-molecule target detection, endows the sensors with inherent selectivity. The structure-switching response is target specific, resulting in minimal effects from nonspecific molecules to the measured current response. Gaining insights into the fundamental mechanisms underpinning aptamer-modified nanopore biosensors expands the versatility of this technology for detecting small molecules. Beyond the development of innovative nanotools that bring us closer to unraveling the complexity of brain chemistry, our findings will drive further innovations in biomimetic nanopore technologies.

## Methods/Experimental

### Materials

Sigma-Aldrich Chemie GmbH (Buchs, Switzerland) was the main supplier for chemicals used in this work, unless otherwise noted. All measurements utilized, phosphate buffer saline (PBS) at 1x concentration (137 mM NaCl, 2.7 mM KCl, 10 mM Na_2_HPO_4_, 1.8 mM KH_2_PO_4_) and pH 7.4 (ThermoFisher Scientific AG, Reinach, Switzerland) as received, or in human serum (Sigma Aldrich). Deionized water (resistivity 18.2 MΩcm^-1^ at 25 °C produced by a Milli-Q system Millipore, Billerica, MA) was used for the preparation of all solutions. All aptamers were purchased and HPLC-purified by Microsynth AG (Balgach, Switzerland). Aptamer stock solutions of 100 *µ*M were aliquoted and stored at −20 °C until use.

The following thiolated single-stranded DNA sequences were used in this work: *dopamine aptamer* ^6^: 5’/Thiol/CGA CGC CAG TTT GAA GGT TCG TTC GCA GGT GTG GAG TGA CGT CG/3’ with molecular weight 13,871.8 g/mol and melting point of 73.7 °C, *scrambled sequence*: 5’/Thiol/ AGT ACG TCG ATG CTC GAT CAG TGG GCT AGG TGC GTA GCG GTC TG/3’ with molecular weight 13,871.8 g/mol and melting point of 71.4 °C, and *serotonin aptamer* ^6^: 5’/Thiol / CGA CTG GTA GGC AGA TAG GGG AAG CTG ATT CGA TGC GTG GGT CG/3’ with molecular weight 13,969.8 g/mol and melting point of 74 °C. The melting temperatures were provided by Microsynth who supplied the DNA sequences.

Dopamine solutions were prepared by mixing dopamine hydrochloride in PBS or in human serum with 10 % wt *L*-ascorbic acid and then serially diluted for the desired concentration. Ascorbic acid was added to decelerate the oxidation of dopamine. ^61,62^

### Nanopipette fabrication and characterization

Nanopipettes were fabricated from quartz capillaries with filament (o.d. 1 mm, i.d. 0.5 mm, 10 cm length, World Precision Instruments QF100-50-10). The capillaries were transformed into nanopipettes using a laser puller (P2000, Sutter Instruments). For reproducible nanopipettes, the laser puller was preheated for at least 1 h prior to use and a pull was activated without fastening a capillary, prior to nanopipette fabrication. To achieve *∼*10 nm diameter orifices, the following parameters were used: (line 1) Heat 750, Filament 4, Velocity 40, Delay 150, and Pull 80; (line 2) Heat 700, Filament 3, Velocity 60, Delay 135, Pull 180. We note that these parameters may vary from instrument-to-instrument^63,64^ and requires fine tuning and characterization with microscopy to confirm nanopore sizes.

### Aptamer functionalization

DNA sequences were functionalized on the inside of the quartz nanopipette using a previously reported protocol.^27^ Briefly, vapor phase deposition was conducted under vacuum at 40 °C for 1 h to assemble monolayers of (3-aminopropyl)trimethoxysilane (APTMS) on the nanopipette surfaces. Maintaining ambient humidity below 40 % is crucial for monolayer assembly, and silane clogging has been observed if this parameter was disregarded. To increase the yield of functional sensors, silanized nanopipettes were first characterized in 1x PBS to ensure proper surface assembly prior to subsequent steps. Then, nanopipettes were filled for 1 h with 1 mM solutions of 3-maleimidobenzoic acid *N*-hydroxysuccinimide ester (MBS) dissolved in a 1:9 (v/v) mixture of dimethyl sulfoxide and PBS. Aptamer disulfide bonds were reduced for 1 h at room temperature using a 50-fold excess of tris(2-carboxyethyl) phosphine (TCEP) relative to DNA aptamer concentration. The DNA solution was diluted to 5 *µ*M in 1x PBS and cleaned to remove unreacted TCEP and cleaved protective groups, using ZebaTM spin desalting columns (7K MWCO, 0.5 mL, ThermoFisher Scientific AG, Reinach, Switzerland). The purified DNA solution was then heated up to 95 °C for 5 min and then renatured by cooling to room temperature to ensure unhybridized sequences in optimal conformations for surface assembly. The MBS solution was removed from the nanopipettes and the sensors were rinsed with 1x PBS. Then, the prepared aptamer solution was incubated for a minimum of 2 h to ensure functionalization to the nanopipette surface. Prior to storage or experimental use, the aptamer solution was removed and the sensors were rinsed three-fold with PBS. Nanopipettes were stored filled with Milli-Q water to reduce etching of the quartz^65^ and were stored at 4 °C in high humidity environments to prevent solution evaporation, which may lead to nanoscale tip breakage upon salt crystal formation. Filling and emptying of the nanopipettes was enabled by MicroFil syringe tips (World Precision Instruments, Sarasota, FL).

### Sensing measurements *via* aptamer-modified nanopipettes

Current measurement was enabled by two Ag/AgCl quasi-reference counter electrodes fabricated in house. One electrode was positioned inside the nanopipette and another in the bulk solution. The current was measured *via* a custom-built high gain current amplifier and the data recorded using a custom written LabVIEW interface (2017, National Instruments), based on WEC-SPM package provided by Warwick Electrochemistry and Interfaces Group. Data was collected using an FPGA card PCIe-7852R (National Instruments). The current magnitudes and potentials reported in the manuscript are denoted with respect to the electrode in the solution bulk. Cyclic voltammograms were acquired by sweeping voltage at 0.2 V s^-1^ voltage sweep rate.

### Quartz crystal microbalance with dissipation monitoring (QCM-D)

The QSense E4 (Biolin Scientific) was used for QCM-D measurements and thiolated aptamers were assembled on QSense gold chips (QSX 301). The chips underwent a rigorous cleaning procedure consisting of 2 min sonication cycles in the following solutions chronologically: isopropanol, acetone, and Milli-Q water. The chips were dried using pressurized nitrogen and then ozone cleaned for 30 min. The QCM-D chip was then sandwiched between two electrodes that apply a voltage to excite the piezoelectric material at its resonance frequency. A winding channel at the flow cell inlet ensures a stable liquid temperature of 24 °C upon contacting the chip surface. The signal from the 3rd harmonic is shown in all graphs. Further details of the measurement protocols and mathematical calculations for extracting the change in monolayer height from the ΔF is available in the Supporting Information.

### Molecular dynamics simulations of aptamer structures

The single-stranded dopamine and serotonin DNA aptamers were first modeled with the Macro Molecule Builder command line tool v2.17.^66^ The final state of these resulting structures were used as a starting point for the MD simulations. The simulation of the dopamine and serotonin aptamers and the interactions with their respective targets were performed in Gromacs using the AMBER99SB-ildn force field. The aptamers were placed in the center of a water box of suitable dimensions according to the size of the different aptamers. Then, the respective analyte, dopamine or serotonin, was inserted into the box, with a 1 Å distance between the box surface and the aptamer. Afterwards, ions and water molecules represented using the transferable intermolecular potential with 3 points (TIP3P) model, were inserted into the box.

Amber Tools in Gromacs was used to generate the analyte topology files. By applying the steepest descent algorithm and considering periodic boundary conditions (PBC) in all directions, energy minimization was performed. The systems were equilibrated in NVT (where the constant number (N), volume (V), and temperature (T), respectively) and NPT (constant number (N), pressure (P), and temperature (T), respectively) and assembled with a time step of 2 fs for 100 ps. Finally, the simulation was performed at room temperature and pressure: 294 K and 1.01 atm for 200 ns. The MD simulation were performed using high performance computing (CINECA). Gromacs tools were used to analyze the trajectories of conformational change (*e.g.*, radius of gyration, RMSD) of the MD simulation. Further details of the RMSD calculations and intermolecular interactions between the aptamer and analyte can be found in the Supporting Information.

### Finite element method simulations

Rectifying behavior of the ionic current through nanopipettes was modeled using finite element method software package Comsol Multiphysics (version 6.0) with Transport of Diluted Species and Electrostatics modules. Further details of the model geometry, equation formulation, mesh, *etc*. are available in the Supporting Information.

### Statistics

All statistics were carried out using GraphPad Prism 9 (GraphPad Software Inc., San Diego). Data are reported as means *±* standard errors of the means with probabilities *P<*0.05 considered statistically significant. Comparative data was evaluated by one-way analysis of variance followed by Tukey’s multiple group comparisons.

## Acknowledgement

The authors thank Prof. Janos Vörös for helpful discussions. A.S. and N. N. thank the ETH Zürich research grant (ETH-13 21-2) and A.D. and D.G. the European Union under the Horizon 2020 Program, FET-Open: DNA-FAIRYLIGHTS, Grant Agreement No. 964995. A.D. also thanks Walter Rocchia for the access to the computation facility. D.M. acknowledges the financial support from the European Research Council (ERC) under the European Union’s Horizon 2020 research and innovation programme (Grant Agreement 948238).

## Supporting Information

The Supporting Information is available free of charge at 20 nm nanopipette characterization (Figure S1); rectification coefficient during sensing (Figure S2); quartz crystal microbalance with dissipation monitoring: protocol, reproducibility, and calculation of monolayer height (Figure S3); root-mean square deviation of molecular dynamic simulations (Figure S4); intermolecular interactions of aptamer binding pocket (Figure S5); finite element modeling of nanopipette sensors (Figures S6-S11, Table 1) (PDF).

This manuscript has been previously submitted to a pre-print server: Annina Stuber, Ali Douaki, Julian Hengsteler, Denis Buckingham, Dmitry Momotenko, Denis Garoli, Nako Nakatsuka. 2023, BioRxiv. https://www.biorxiv.org/content/10.1101/2023.03.10.532011v1 (August 31, 2023).

---

**TOC Graphic.**
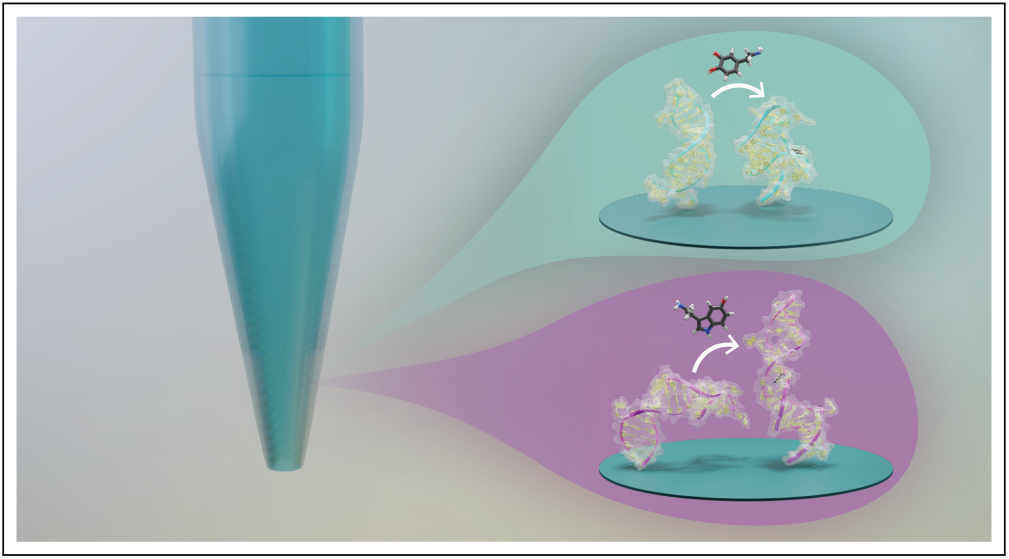

