## Supporting Information for "Aptamer Conformational Dynamics Modulate Neurotransmitter Sensing in Nanopores"

### Contents

**SI-1. 20 nm diameter nanopipette sensing** . . . . . . . . . . . . . . . . . . . . **S3**

**SI-2. Rectification coefficient during sensing** . . . . . . . . . . . . . . . . . . . **S5**

**SI-3. Quartz crystal microbalance with dissipation monitoring** . . . . . . . . **S6**

Measurement protocol . . . . . . . . . . . . . . . . . . . . . . . . . . . . . . . . S6

Calculating monolayer height *via* the Sauerbrey equation . . . . . . . . . . . . . . S7

**SI-4. Molecular dynamic simulations (MD simulations)** . . . . . . . . . . . . **S9**

Root-mean square deviation (RMSD) . . . . . . . . . . . . . . . . . . . . . . . . S9

Intermolecular interactions in the aptamer binding pocket . . . . . . . . . . . . S10

**SI-5. Modeling rectifying behavior of aptamer-modified nanopipettes** . . . . **S11**

Model description . . . . . . . . . . . . . . . . . . . . . . . . . . . . . . . . . S11

Estimation of the nanopore opening . . . . . . . . . . . . . . . . . . . . . . . . S14

Aptamer-modified nanopore sensor optimization: effect of the pore size . . . . . . S15

Effect of charged aptamer layer on measured current through nanopipettes . . . . S16

Aptamer layer compression that incorporates change of charge density . . . . . . . S17

Aptamer layer compression that incorporates ion permeability . . . . . . . . . . . S18

Aptamer-modified nanopore sensor optimization: effect of the aptamer layer thickness and its variation . . . . . . . . . . . . . . . . . . . . . . . . . . . . . S19

**SI-6. Dukhin number calculation** . . . . . . . . . . . . . . . . . . . . . . . . . **S21**

**SI-7. Estimation of theoretical current** . . . . . . . . . . . . . . . . . . . . . **S22**

### SI-1. 20 nm diameter nanopipette sensing

Nanopipettes with 20 nm diameter were fabricated from the same quartz capillaries as used for the 10 nm diameter nanopipettes, (o.d. 1 mm, i.d. 0.5 mm, 10 cm length, World Precision Instruments QF100-50-10). The laser puller (P2000, Sutter Instruments) was used with the following program: (line 1) Heat 850, Filament 4, Velocity 30, Delay 145, and Pull 110; (line 2) Heat 865, Filament 3, Velocity 20, Delay 135, Pull 180. The same protocol was followed as described in the main text for the 10 nm diameter nanopipettes and the process was monitored using ion current rectification (Figure S1a).

While the characteristics of the current-voltage curves are retained *vs.* 10 nm nanopores indicating successful functionalization (main text, Figure 1a, b), the current range is widened due to the larger pore size. Upon exposure of the aptamer-modified 20 nm nanopipettes to 100 uM dopamine, negligible changes in the current response were observed. The schematic in Figure S1b shows how based on the size of the dopamine aptamers (extracted from MD simulations), the structure-switching dynamics of the aptamers is negligible inside of the 20 nm pore. Thus, the ionic current is minimally altered upon dopamine recognition (despite addition of +1 charge due to dopamine), indicating the importance of the nanopore dimension to harness the aptamer switch for signal transduction.

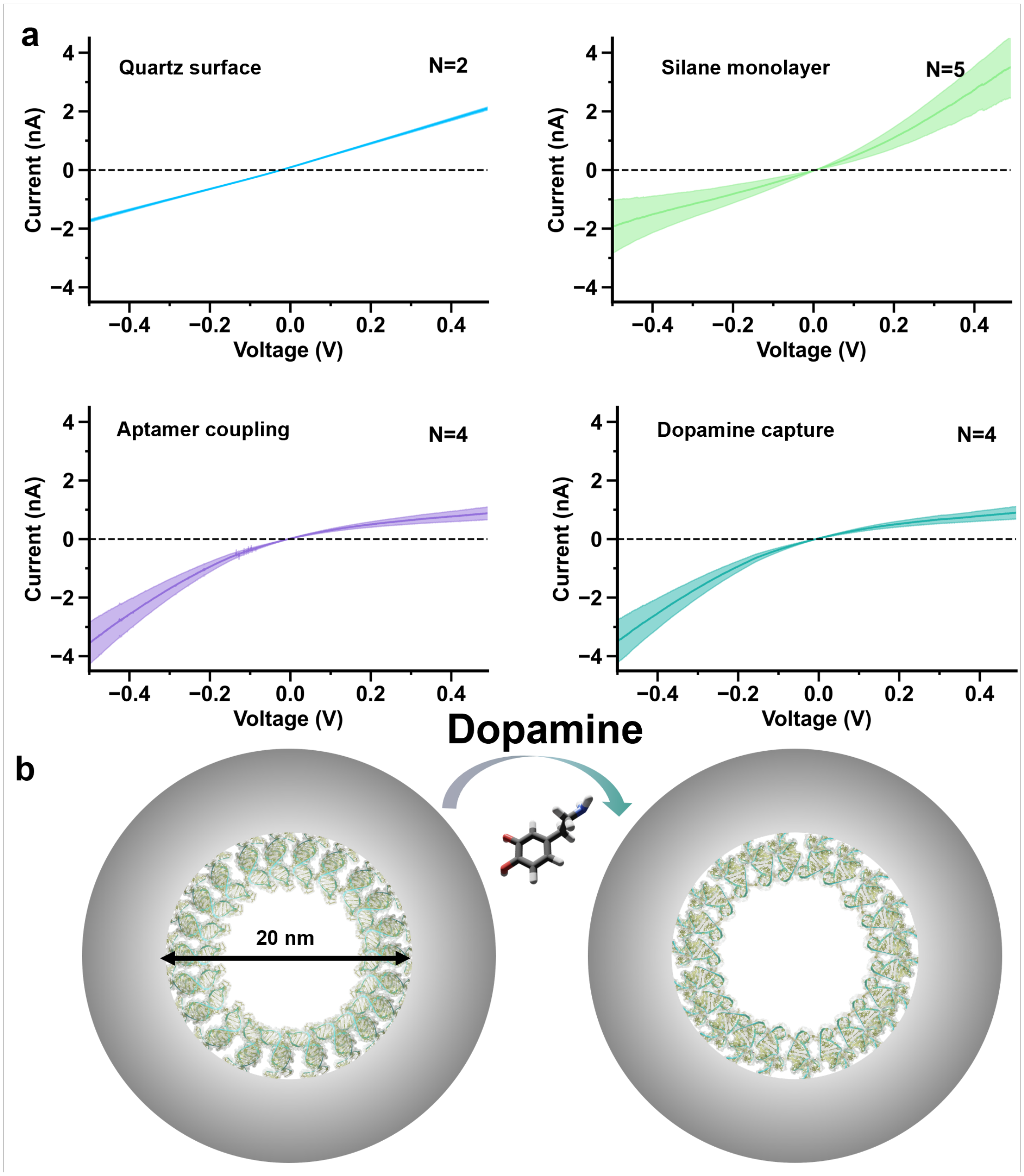


Figure. S1: (a) Functionalization process of aptamers through sequential surface chemistry can be monitored using the ion current rectification (ICR) effect manifested as asymmetric current vs. voltage curves. Negative charges on bare quartz nanopipettes ($N$=2) are inverted to positive upon assembly of aminosilanes ($N$=5). Coupling of negatively charged aptamers leads to higher rectification behavior ($N$=4) followed by exposure of the 20 nm diameter nanopipette to 100 µM dopamine with 10 % weight ascorbic acid. The solid line represents the average and the shaded area the standard error of the mean.(b) Schematic view inside of a 20 nm orifice nanopipette lined with aptamers pre- (left) and post- (right) dopamine exposure.

#### SI-2. Rectification coefficient during sensing

We extracted the rectification coefficient ($r$) from the current-voltage (I-V) curves, a value that enables direct comparisons of surface charge at the walls of the nanopipette.[1] As aptamer conformational changes rearrange negative charge densities at the surfaces to which they are covalently bound, we expected the $r$ value to be influenced in a target-specific manner. This coefficient is calculated by taking the logarithm of the ratio of the absolute values of the current measured at a positively applied voltage, by the current at the corresponding negative voltage:

$$r = log(|I_{V+}/I_{V-}|)$$

We observed opposite directionalities of $r$ of the serotonin and dopamine nanopipettes upon their respective analyte exposures (Figure S2). For the dopamine aptamer-modified sensor, the $r$ value becomes increasingly negative, from -0.5 ± 0.03 to -0.6 ± 0.05 upon exposure to dopamine. Conversely, the $r$ value of the serotonin aptamer-modified sensor increases in value upon serotonin capture from -0.7 ± 0.03 to -0.4 ± 0.06.

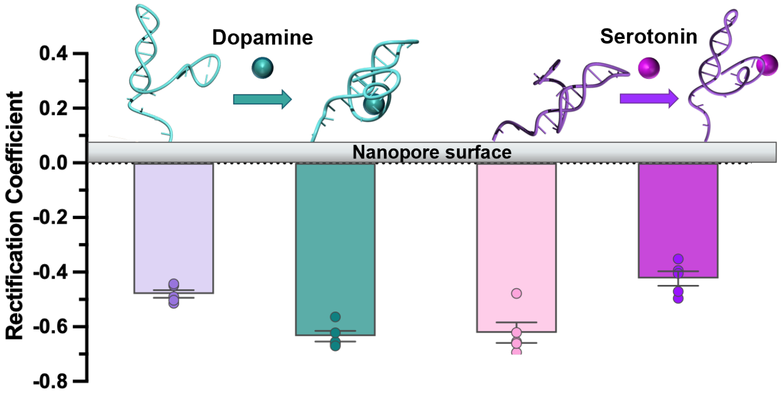


Figure. S2: The rectification coefficient $r$ was calculated from the ion current rectification (ICR) plots at +0.5 V and -0.5 V. For dopamine aptamers, the $r$ value becomes more negative upon dopamine capture [$F(8)=3.877$, $P<0.05$]. In contrast, for serotonin aptamers, the $r$ value becomes less negative upon serotonin recognition [$F(8)=4.301$, $P<0.01$]. Error bars represent standard errors of the mean for $N=5$ sensors.

#### SI-3. Quartz crystal microbalance with dissipation monitoring

##### Measurement protocol

A stable baseline was obtained in PBS prior to aptamer assembly. The thiolated DNA sequences were pre-treated following the same procedure as the nanopipette functionalization (cleaving, dilution, purification, heat treatment). The aptamer solution was injected and incubated until the frequency (Figure 3a) and dissipation (Figure S3a) plateaued. Then, covalent binding of the aptamer was challenged by flushing buffer to rinse any unbound sequences. Dopamine prepared with 10 wt % ascorbic acid was exposed to the chip and was then washed away by flushing the buffer. The same surface was used for multiple repeats of dopamine binding and unbinding (Figure S3b). Repeated measures were conducted on $N$=3 different QCM-D chips for a total of $N$=6 measurements including retesting on certain chips (Figure S3c).

For certain repeated measures, the aptamer solution was incubated overnight on the QCM-D chips rather than assembled *in situ*; both functionalization routes yielded comparable magnitude responses to dopamine. The same procedure was followed using scrambled DNA-modified chips to ensure sequence-specific changes in frequency and dissipation (Figure S3d). We investigated the frequency change upon exposing the dopamine aptamer-modified surface to increasing concentrations of dopamine. (Figure S3e,d) The frequency increased with increasing amounts of dopamine and saturated in the $\mu$M range. However, this detection range and sensitivity is not comparable to interactions inside nanopipettes. On the QCM-D chip, we measure the combined effect of added small-molecule mass and aptamer conformational changes that alter adsorption of water molecules to the surface. Within the nanoscale orifice, we do not rely on analyte mass, but solely on the structure-switching aptamers to modulate the current response.

#### Calculating monolayer height *via* the Sauerbrey equation

The change in mass on the quartz crystal can be calculated from a measured change in frequency ($\Delta(f)$) at a specific harmonic ($n$) as described by the Sauerbrey equation:

$$m_{QCM} = -c\Delta f/n \quad (1)$$

where the mass sensitivity constant, $c = 17.7$ ng/cm$^2$Hz relates to the properties for the 5 MHz quartz crystal chip. The height or the effective distance from the chip surface ($d_{eff}$) can then be extracted using the equation:

$$d_{eff} = \Delta m/\rho_{\mathrm{H_2O}} \quad (2)$$

where $\Delta m$ is the change in mass on the chip calculated prior, and $\rho_{\mathrm{H_2O}}$ the density of water molecules displaced upon aptamer assembly or structure switching. This direct relation, linking change of frequency to change in height, is only applicable for quasi-rigid films, which have the required condition that $\Delta D/\Delta f < 4{*}10^{-7}$ Hz$^{-1}$.[2,3] Our layer is classified as quasi-rigid as the $\Delta D/\Delta f$ of our aptamers assembled on the chip was calculated to be $\sim 1 * 10^{-7}$ Hz$^{-1}$.

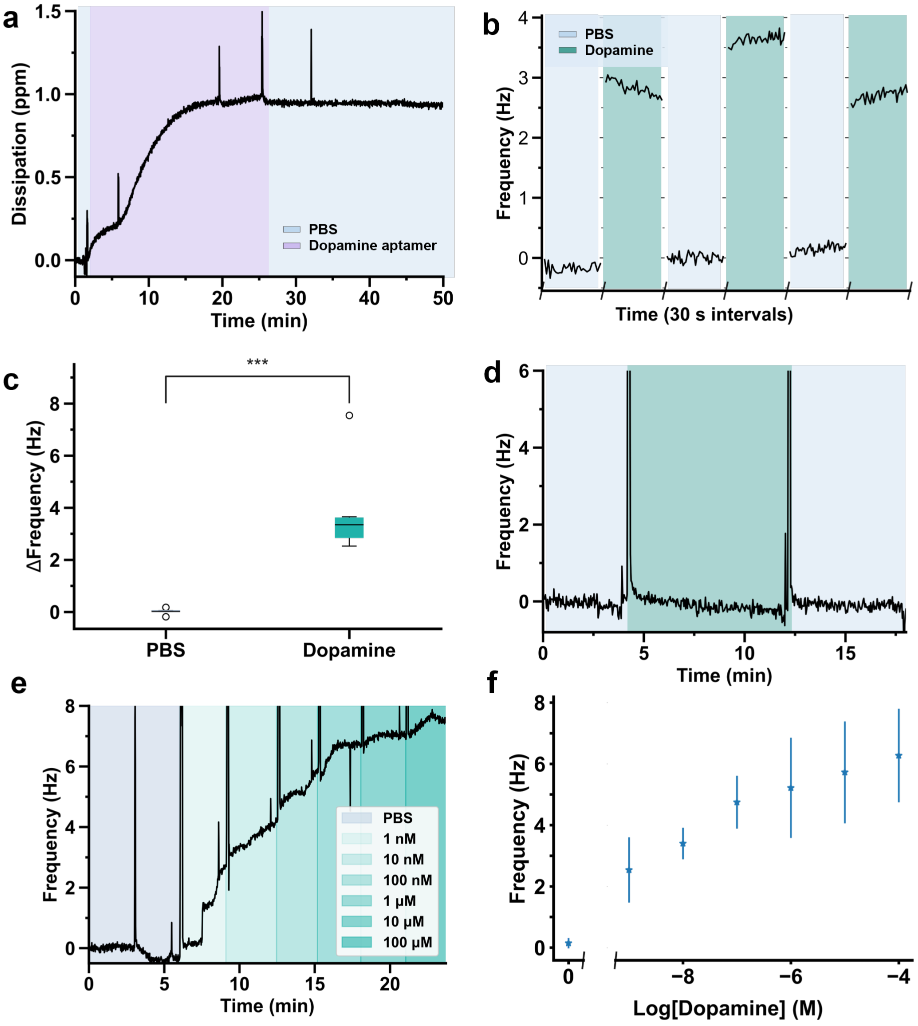


Figure. S3: (a) Dopamine aptamer assembly in phosphate buffered saline (PBS) results in a dissipation increase of ∼1 parts per million (ppm). (b) Three consecutive repeats of switching the solution on a single chip from PBS to dopamine solution demonstrating the reproducibility of the reversible signal due to binding and unbinding of dopamine from the aptamer. (c) The frequency change upon PBS injection *vs.* dopamine injection results in a statistically significant difference ($P<0.001$) measured on three different QCM-D sensors for a total of $N=6$ repeated measures. (d) Control measurements run in parallel where 100 $\mu$M of dopamine solution is exposed to a QCM-D sensor functionalized with the scrambled DNA sequence (same nucleotides as the specific dopamine aptamer, but in a random order to hinder dopamine recognition), resulted in negiligible changes in the frequency. (e) Various concentrations of dopamine prepared with 10% weight ascorbic acid, were washed over a aptamer-functionalized QCM chip. (f) The average frequency was extracted over the time period of concentration specific exposure. Two chips *(N=2)* were exposed to different concentrations in parallel, and the average of the two points with their respective standard deviations are represented on a logarithmic scale. For all measurements, the 3rd overtone is shown, and the baseline is set to 0 Hz.

#### SI-4. Molecular dynamic (MD) simulations

##### Root-mean square deviation (RMSD)

The following figure illustrates the change in the RMSD values of dopamine and serotonin aptamers after 100 ns of simulations (Figure S4). The RMSD values of the dopamine and serotonin aptamers were 0.7 nm and 1.2 nm, respectively. These low values show that 100 ns of simulation was enough to reach a stable state, hence, the final molecular structures shown in the main text (Figure 3) were performed for 100 ns.

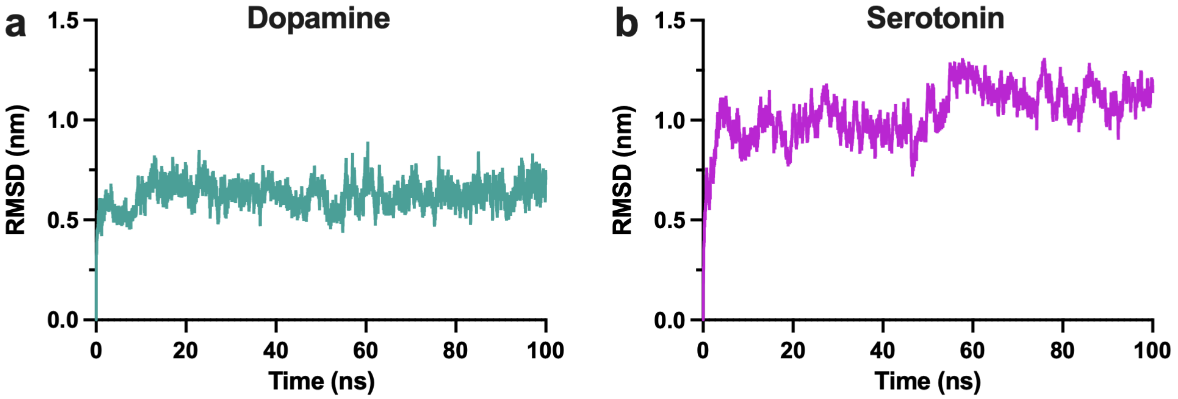


Figure. S4: Root-mean square deviation values of (a) dopamine and (b) serotonin aptamers.

#### Intermolecular interactions in the aptamer binding pocket

At the end of the 100 ns simulation, the intermolecular interactions between the dopamine aptamer (Figure S3a) and the serotonin aptamer (Figure S3b) and their respective targets were extracted in 2-D.

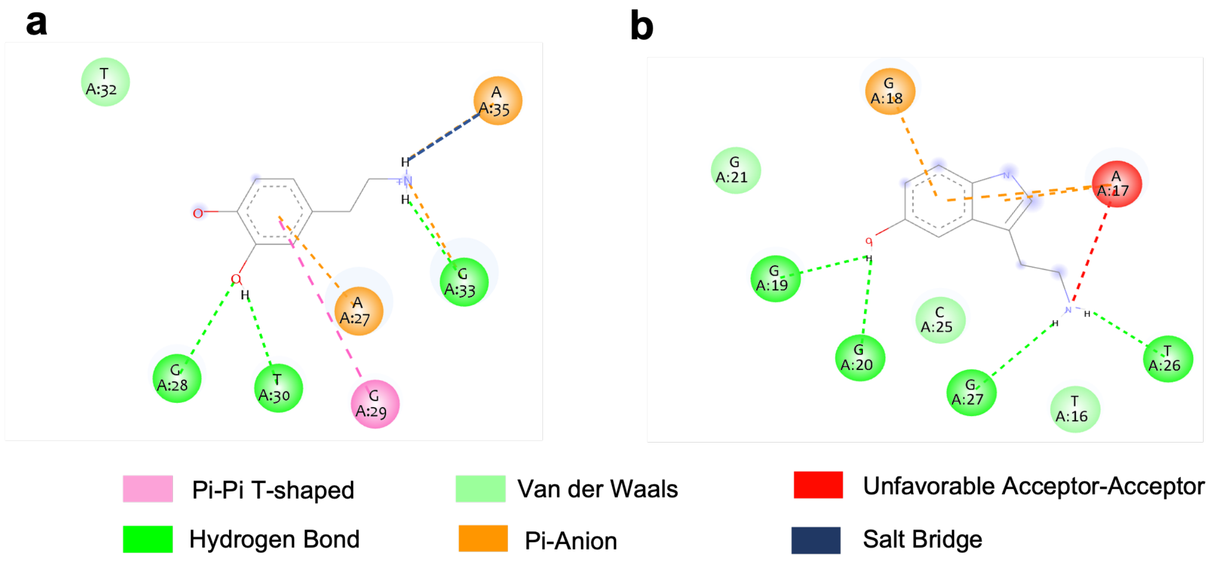


Figure. S5: Interactions established in two dimensions extracted at the end of the 100 ns molecular dynamic simulation for (a) the dopamine aptamer-dopamine interactions and (b) the serotonin aptamer-serotonin interactions. Aptamer nucleotides are represented by spheres of different colours around the structure. Green dashed lines represents the H-bonds, pink the Pi-Pi T-shaped interactions, orange the Pi-anion interactions, red the unfavorable acceptor-acceptor interactions, and blue the salt bridge. The light green nucleotides void of dotted line links, are involved in Van der Waals interactions with the analyte.

#### SI-5. Modeling rectifying behavior of aptamer-modified nanopipettes

##### Model description

The finite element model, including the geometry, mesh, and equation formulations is based on our previous work for serotonin-specific aptamer-modified nanopore sensors.[4] The model allows the simulation of ion transport in the presence of an electrical double layer near the pipette walls. Briefly, the simulations domain is built by taking advantage of the geometry's axial symmetry, allowing reduction of computational efforts. The geometry of the computational domain (10 µm x 2 µm) includes a 7.5 µm long section of the nanopipette with appropriate inner and outer half-cone angles, wall thickness at the tip, and an opening radius of 4.6 nm, in accordance with the data from TEM images.[4] The model is discretized into *ca.* 680,000 triangular mesh elements with minimum element size of 0.1 nm. The size of the domain and the nanopipette are optimized to have minimal influence on the numerical result.

The numerical model is formulated using Nernst-Planck:

$$J_i = -D_i \nabla c_i - (z_i F)/RT D_i(T) c_i \nabla \varphi \tag{3}$$

and Poisson equations:

$$\nabla^2 \varphi = -F/(\epsilon \epsilon_0) \Sigma (z_i c_i) \tag{4}$$

that describe ion transport (in the absence of liquid flow) as well as the existence of accumulation of charged species in solution, respectively (Poisson equation allows the electroneutrality to be broken). Here, $J_i$, $D_i$, $z_i$, and $c_i$, denote the flux, diffusion coefficient, charge number, and local concentration of ionic species $i$, while $\varphi$, $F$, $R$, $T$, $\epsilon$ and $\epsilon_0$ specify the electrical potential, Faraday and gas constants, temperature, dielectric constant, and vacuum permittivity, respectively. The model is resolved in the geometry of a nanopipette using appropriate boundary conditions as shown in Figure S6 with boundary conditions specified in Table 1. The thickness of the aptamer layer was varied from 5.5 nm (simulating a case with no analyte present) to 4.9 nm (in presence of dopamine), based on values extracted from the molecular dynamic (MD) simulations.

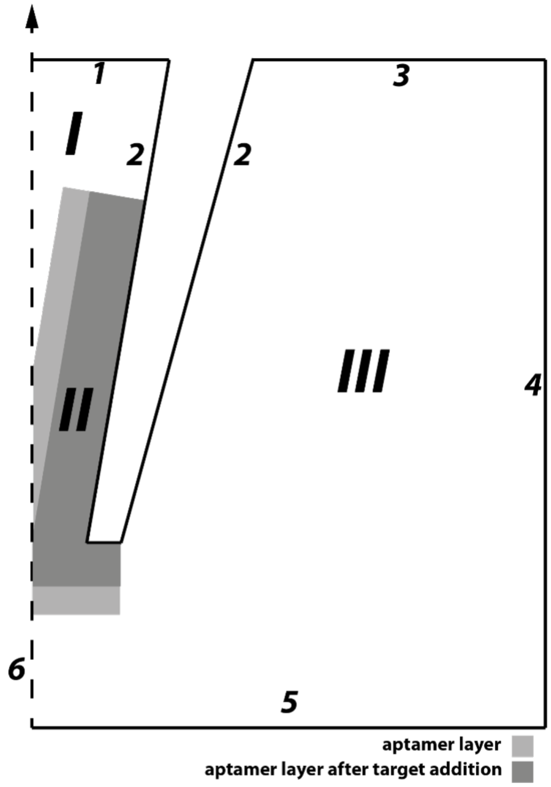


Figure. S6: Schematic representation of the model geometry. Boundary and domain conditions are specified in Table 1 in accordance with the designation on the graph. The change of the aptamer layer (domain II) thickness is illustrated with a grey color shade.

The boundary conditions define electrodes with applied potential and determine bulk solution properties. In particular, the diffusion coefficient inside the aptamer layer (domain II, grey shading in Figure S6) determines the mobility of ions through the layer and is modeled simply as the product of the bulk diffusion constant multiplied by a factor $f \leq 1$. The charge density of the aptamer layer also differs from the bulk electrolyte as the aptamer backbone is highly negatively charged. Individual aptamer charges are taken into account by varying the concentration of these fixed charges in the domain II, while charge density in the bulk solution (domains I and III) is determined only by mobile ions $Na^+$ and $Cl^-$.

Table 1: Boundary conditions of the model and specific parameters in model domains. Symbol $n$ denotes the vector normal to the boundary.

| **Boundary** | **Nernst-Planck equation** | **Poisson equation** |
|---|---|---|
| 1 | concentration, $c_i$ = 158 mM | applied potential, $\varphi$= bias |
| 2 | no flux, -$n \cdot J_j$=0 | no charge, $n \cdot \nabla \phi = 0$ |
| 3 | no flux, -$n \cdot J_j$=0 | surface charge, $n \cdot \nabla \phi = \sigma$ |
| 4 | concentration, $c_i$=158 mM | ground, $\varphi$= 0 |
| 5 | no flux, -n$\cdot J_j$=0 | no charge, $n \cdot \nabla \phi = 0$ |
| 6 | axial symmetry | axial symmetry |
| **Domain** | **Nernst-Planck equation** | **Poisson equation** |
| I | diffusion coefficient, $D_i$ | charge density, $\rho = c_{Na^+} - c_{Cl^-}$ |
| II | diffusion coefficient, $fD_i$ | charge density, $\rho = c_{Na^+} - c_{Cl^-} - c_{fixed}$ |
| III | diffusion coefficient, $D_i$ | charge density, $\rho = c_{Na^+} - c_{Cl^-}$ |

**Estimation of the nanopore opening**

To obtain the correct value of the nanopipette opening, the experimental electrical resistance of the bare nanopipette was matched with the simulation results at different opening radii. As the experimental $I - V$ curves reveal significant rectifying behaviour, only a linear part of the experimental voltammogram (between -10 and +10 mV) was used for this purpose. A linear fit of the $I - V$ curve indicated a resistance value of 367 MOhm, which matches numerical result for a pipette with opening diameter 9.2 nm.

**Aptamer-modified nanopore sensor optimization: effect of the pore size**

For the fixed aptamer configuration (5.5 nm/4.9 nm thick layer in the absence/presence of analyte), the model also allows to find optimal nanopore size. As shown in Figure S7, the pore reveals maximal response upon analyte binding for the opening diameter of 8.4 nm. At smaller pore sizes, there is a slight decrease of the pipette response towards analyte addition, while for nanopores with apertures >10 nm the response drops very quickly. Thus, larger pores are unlikely to be suitable for sensing. For the dopamine-sensitive aptamer employed in the current work, the response is very close to the optimum (*ca.* 99 % of the optimal current response at 8.4 nm).

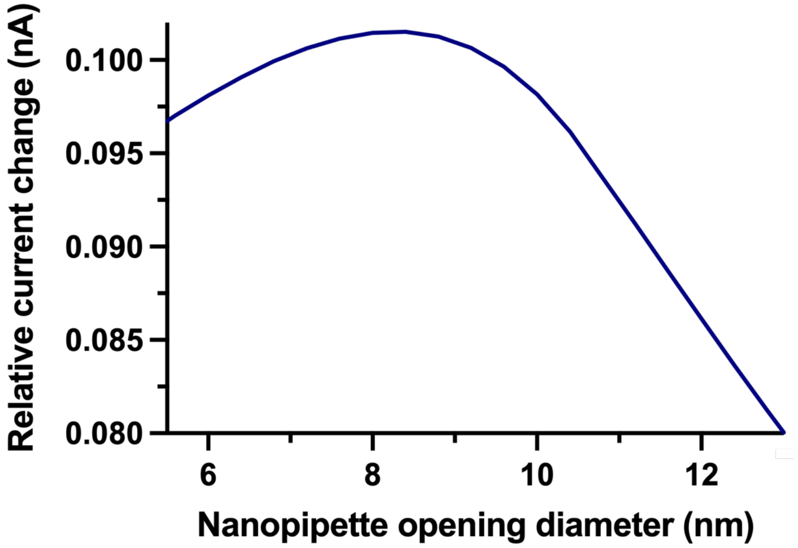


Figure. S7: Relative current change upon aptamer layer thickness variation from 5.5 nm to 13 nm for nanopores with different opening sizes.

#### Effect of charged aptamer layer on measured current through nanopipettes

The effect of the fixed charges inside the aptamer layer was then modeled for the nanopipette sensors modified with dopamine aptamers prior to exposure to the target (domain II with aptamer layer thickness of 5.5 nm, light gray shading in Figure S6) and post dopamine binding (domain II with aptamer layer thickness of 4.9 nm, dark gray shading in Figure S6). As can be seen from Figure S8, the presence of the fixed charges in the aptamer layer leads to an increase in the ion current rectification when the layer gets progressively more negatively charged (from 0 to 200 mM of charges). Under these assumptions, the thickness of the aptamer layer plays little role in the magnitude of the ionic current - solid (5.5 nm) and dotted (4.9 nm) lines almost coincide.

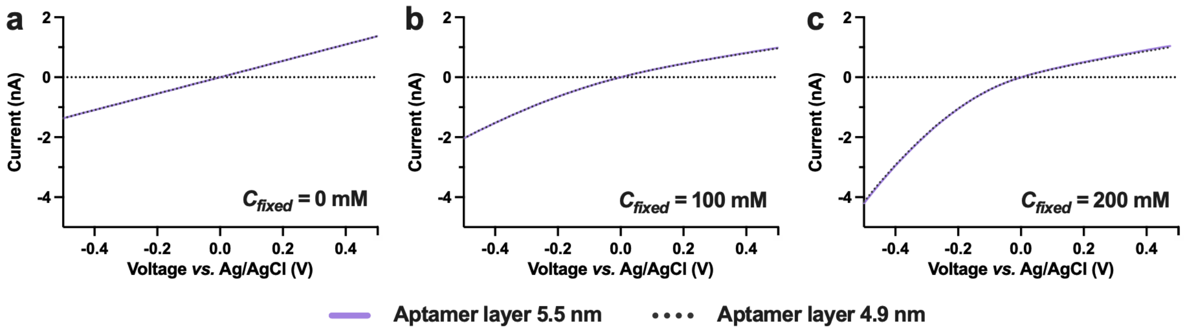


Figure. S8: Simulated $I-V$ characteristics of the nanopipettes with aptamer layer thicknesses of 5.5 (solid purple line) and 4.9 nm (dotted grey line) and concentration of fixed charges within the layer of (a) 0 mM, (b) 100 mM and (c) 200 mM.

**Aptamer layer compression that incorporates change of charge density**

To understand the effect of aptamer layer compression on the ionic current beyond changes in the layer height, the model for 4.9 nm-thick aptamer layer incorporated a proportional increase of the charge density by 12%. This value stems from the height ratio between the extended *vs.* contracted aptamer molecule (5.5 *vs.* 4.9 nm). The increase in the charge density upon target recognition, manifests as an increasingly pronounced rectification, with higher current magnitudes at negative bias and slightly lower currents in the positive $I-V$ curve branch (Figure S9). Yet, there is still a negligible difference between the free *vs.* target-bound aptamer layer curves at positive voltages.

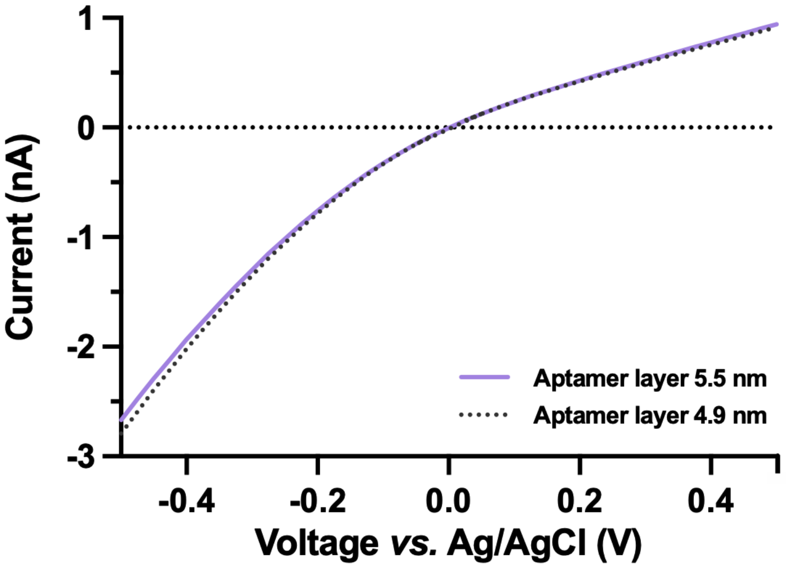


Figure. S9: Simulated $I-V$ characteristics of the nanopipettes with aptamer layer thicknesses of 5.5 (solid purple line) and 4.9 nm (dotted grey line) with concentration of fixed charges of 200 mM and 224.5 mM ((5.5 nm/4.9 nm)*200 mM) respectively.

##### Aptamer layer compression that incorporates ion permeability

In addition to the effect of charge density changes upon target capture, the change of ion permeability through the aptamer layer was also introduced into the model by varying the factor $f$ from 1 to 4.9 nm/5.5 nm = 0.891. Changing this value affects the diffusion constant $D_i$ of all ionic species in the domain II of the model. Figure S10 shows how the ion mobility in the aptamer layer impacts the overall ionic current in the aptamer-modified nanopore. In this model, the ion current magnitude is visibly smaller at positive biases upon compressing the aptamer layer and lowering the ion mobility (solid purple curve *vs.* red dashed curve). Negligible differences are observed at negative voltages upon compressing the layer and reducing the ion mobility as the effects of altered ion mobility and more compact charge have opposite effects. The overlapping effects of reduced permeability hindering ionic flux and higher density of charge increasing the flux, counteract each effect, causing a less pronounced ion current change.

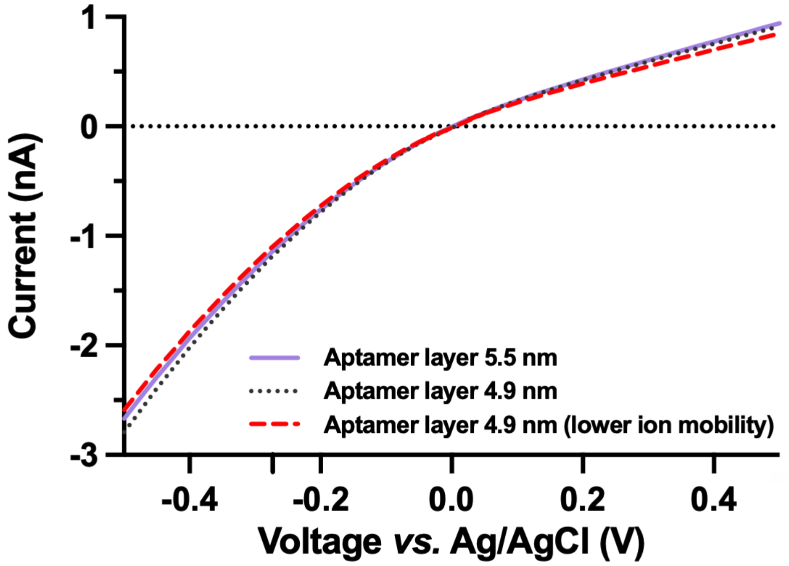


Figure. S10: Simulated $I-V$ characteristics of the nanopipettes with aptamer layer thickness of 5.5 (solid purple line) and 4.9 nm (dotted grey line), with concentration of fixed charges of 200 mM and 5.5/4.9 * 200 mM = 224.5 mM, respectively. The dashed red line reveals ion currents simulated for the contracted aptamer layer with proportionally reduced ion mobility.

**Aptamer-modified nanopore sensor optimization: effect of the aptamer layer thickness and its variation**

The simulations shown above highlight the need to incorporate the variation of the ionic diffusivity (via parameter f), and the charge concentration in the aptamer layer upon analyte recognition. Taking these factors into account, we modeled ionic current through a nanopipette at different aptamer layer thicknesses, assuming that a 5.5 nm thick aptamer layer contains 200 mM of uncompensated charge and the diffusivity in this case is the same as in the solution bulk (f = 1). For all other cases, the change of the charge concentration, $c_{apt}$, is proportional to the thickness, $\delta$, relative to the 5.5 nm initial aptamer layer thickness value:

$$c_{apt} = \frac{5.5nm}{\delta} \tag{5}$$

whereas diffusivity variation is the inverse:

$$f = \frac{\delta}{5.5nm} \tag{6}$$

Taking these factors into account, we obtained the current response through the nanopore as the function of aptamer layer thickness as shown in Figure S11a. This curve can be easily transformed for establishing a variation of the relative current change upon shrinkage of the aptamer layer by 0.6 nm (similarly to the variation of 5.5 to 4.9 nm but for different layer thicknesses), simulating target recognition (Figure S11b). The maximal response is observed for 5.0 nm aptamer layer thickness, highlighting that our aptamer-nanopore system is almost fully optimal (ca. 98 % of that maximal current variation). We also demonstrate how aptamers that undergo larger magnitudes of conformational change, in this case shown as a greater aptamer layer shrinkage upon dopamine recognition, the relative current change is greater (Figure S11c).

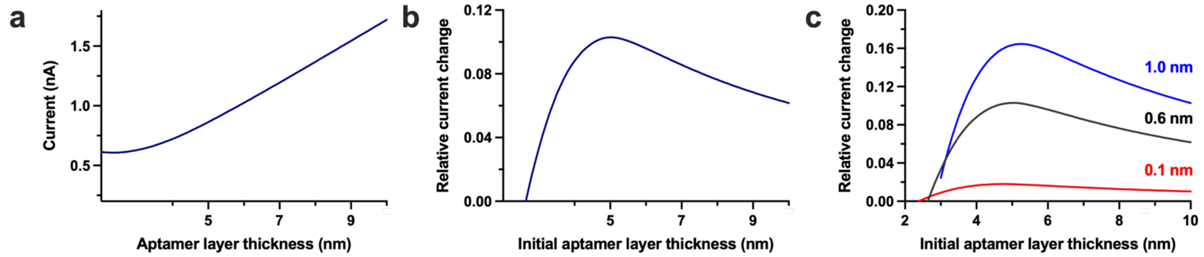


Figure. S11: a) Variation of the current through the nanopore at different aptamer layer thicknesses at 0.5 V bias for a pore with the opening of 9.2 nm. b) Relative current change for the aptamer-modified nanopore with different aptamer layer thicknesses assuming 0.6 nm layer shrinkage (predicted from molecular dynamic simulations) upon target recognition. (c) Relative current change curves for aptamer-modified nanopores with different initial aptamer layer thicknesses calculated for different thickness variations (0.1 nm, 0.6 nm, and 1.0 nm for red, black, and blue traces, respectively).

#### SI-6. Dukhin number calculation

The Dukhin number ($Du$) was approximated to compare the ratio of ionic flux occurring along the nanopipette walls *vs.* through the center of the nanopipette:[5]

$$Du = (|\sigma|/e)/c * R \tag{7}$$

Where $e$ represents the elementary charge and $c$ the ionic bulk concentration at 150 mM, corresponding to the salt content of 1x PBS. A value of 5 nm was used for the nanopore radius, $R$. The surface charge density, $\sigma$, was approximated by projecting the number of negative charges associated with one aptamer (44 negative charges per sequence), onto the surface required for the unbound state of an aptamer; the lower rectangular surface from the tightest fitting box, extracted from simulations of the respective aptamer (Figure 4).

Our approximations result in a Dukhin number of around 8.4 and 4.0 for the dopamine and serotonin aptamers tethered within a 10 nm nanopore, respectively. Both of these values imply that the contribution of the overall ionic flux originating from the flux along the nanopore wall dominate when compared to bulk flux. We note however, that the surface charge density approximation disregards the height of the aptamer, which covers a large portion of the nanoscale orifice. Therefore, a simple projection onto the wall does not necessarily represent the true wall charge of the nanopore.

### SI-7. Estimation of theoretical current

To estimate the current passing through the area of the nanopore covered with DNA, the formula from Wang et al.[6] was used. Here the experimentally obtained signal, $I_{measured}$, can be described as:

$$I_{measured} = I_{open} - I_{excluded} + I_{aptamer} \tag{8}$$

$I_{open}$: current through the bare unmodified nanopore (value obtained experimentally).

$I_{excluded}$: current excluded due to the presence of aptamers immobilized inside the nanopore.

$I_{aptamer}$: current passing through functionalized layer of aptamers.

By utilizing the dimensions extracted from the MD simulations, the $I_{excluded}$ was estimated from the excluded volume:[6]

$$I_{excluded} = \frac{V}{L_{pore}}(N * area_{aptamer}(\mu_{Na} + \mu_{Cl})cN_A e) \tag{9}$$

where $V$ is the applied voltage (500 mV), $L_{pore}$ is the sensing length of the nanopore (simulated to be ~30 nm),[7] $\mu_{Na}$ and $\mu_{Cl}$ are the bulk mobilites of $Na^+$ and $Cl^-$ ions ($5.18 * 10^{-8}$ $m^2$ $V^{-1}$ $s^{-1}$ and $7.91 * 10^{-8}$ $m^2$ $V^{-1}$ $s^{-1}$, respectively[8]), $c$ is the molar concentration of NaCl in mol/$m^3$, $N_A$ is Avogadro's number, and $e$ is the fundamental charge.[9] The $area_{aptamer}$ represents the cross-sectional area occupied by one aptamer within the orifice. This value was calculated from the dimensions of the aptamers extracted from the MD simulations (dimensions are found in the main text, Figure 3, 3.95*5.46=21.56 $nm^2$ and 4.21*5.72=24.08 $nm^2$ for dopamine and serotonin aptamers, respectively). The $N$ represents the number of aptamers within the orifice, which can be estimated as the number of aptamers capable of binding within the 10 nm diameter opening, by dividing the circumference by the dimensions of the lower rectangle from the simulated aptamer structures (Figure 3c,f).

According to equation 9, the excluded current changes upon introduction of the neurotransmitter target for both the dopamine and serotonin aptamers as the cross-sectional area of DNA through which ionic flux traverses is modified upon conformational rearrangement.

A compression of the aptamer layer, as is the case upon dopamine binding, results in a lower projected area of DNA coverage within the pore, and thus reduces the excluded current, while the inverse can be approximated for an aptamer layer elongation.

The $I_{excluded}$ values for dopamine and serotonin were calculated as relative values before and after target exposure, and opposite directionalities of current change were observed. Specifically, $I_{excluded}$ *decreased* by around 20% for dopamine and *increased* by approximately 10% for serotonin. Subsequently, $I_{aptamer}$ was obtained according to equation 6, resulting in the same exhibited directionality of behavior as $I_{excluded}$ for both dopamine and serotonin.

It is however important to note that these values, $I_{excluded}$ and $I_{aptamer}$, depend on the parameters used in equation 7, in particular, $N$, $L_{pore}$, and $c$. Nevertheless, fixing the same values for dopamine and serotonin, the divergent behaviour is always observed (*ie.* a decreasing current for dopamine and an increasing current for serotonin). These directional changes in ionic currents support both the experimental observations and COMSOL simulations: when the aptamer layers are compressed, a decrease in ionic flux is observed upon dopamine binding. Conversely, when the aptamer layer is elongated due to serotonin binding, an increase in ionic flux is reported.
